# IFN-κ directs antiviral immunity in human skin

**DOI:** 10.64898/2026.09.19.752886

**Authors:** Jafira M. Johnson, Laurellee Payne, Pooja Parameswaran, Yongzhi Chen, Mrinal K. Sarkar, Anthony M. Coon, J. Michelle Kahlenberg, Johann E. Gudjonsson, Katherine A. Fitzgerald, Mehdi Rashighi, Megan H. Orzalli

## Abstract

Inducible expression of type I IFNs is a well-established host innate defense mechanism to limit virus infection. Yet, many viruses have evolved strategies to suppress the induction of these cytokines to enhance replication, spread, and transmission between hosts. Whether additional antiviral mechanisms protect against infection when inducible responses are compromised is not well understood. Here, we demonstrate that human keratinocytes preemptively protect the skin against virus infection through constitutive production of IFN-kappa (IFN-κ), a poorly studied type I IFN family member. We find that constitutive *IFNK* expression protects keratinocytes against skin tropic RNA and DNA viruses, including vesicular stomatitis virus (VSV) and herpes simplex virus-1 (HSV-1). Using a human skin organoid model, we further demonstrate that keratinocyte-derived IFN-κ establishes an antiviral state in dermal fibroblasts. Genetic and chemical analysis of the type I IFN receptor (IFNAR) signaling pathway in monocultured keratinocytes and skin organoids revealed that distinct pathways control VSV and HSV-1 replication. While canonical JAK-STAT signaling provided protection against VSV, HSV-1 infection was controlled through a JAK-STAT-independent mechanism. Using transcriptomic analysis, we further identified an IFN-κ-dependent gene signature in keratinocytes that is not similarly driven by other type I IFNs. Together, this work establishes constitutive IFN-κ production by keratinocytes as a broadly antiviral tissue autonomous defense mechanism.

## Introduction

A core principle of innate antiviral immunity is the ability of the host to induce an antiviral transcriptional response following viral infection. Classically, this response is initiated when pathogen-associated molecular patterns (PAMPs) are detected by intracellular pattern recognition receptors (PRRs). Upon activation, PRRs engage downstream signaling pathways that culminate in the activation of transcription factors, such as IRF3/7, NF-κB, and AP1. One of the best-characterized transcriptional outcomes of PRR engagement is the inducible expression of type I interferons (IFNs), a family of cytokines with potent antiviral and antiproliferative properties (*1–3*).

In humans, 17 type I IFN family members have been described, including 13 IFN-*a* subtypes, IFN-β, IFN-ε, IFN-κ, and IFN-ω, with IFN-*a* and IFN-β being the best studied. These cytokines can be secreted by infected cells and signal in an autocrine or paracrine manner through the type I IFN receptor (IFNAR), composed of IFNAR1 and IFNAR2, to activate JAK-STAT signaling and induce the expression of IFN-stimulated genes (ISGs)(*1–3*). The protein products of these ISGs can restrict virus infection by targeting multiple stages of the viral replication cycle, including entry, replication, assembly, and egress, and can also enhance viral sensing or recruit additional immune cells to the site of infection (*3, 4*).

Although inducible production of type I IFN cytokines can be highly effective in controlling some viruses, many human pathogens encode gene products that inhibit either the induction or signaling capacity of these antiviral cytokines, thereby evading this host-defense strategy (*5–7*). When inducible antiviral responses are suppressed, the host may instead rely on constitutively expressed antiviral mechanisms that are present prior to infection to limit viral replication. Such pre-existing defenses may be particularly important at initial sites of infection, such as barrier tissues, where early control of viral replication is critical for limiting subsequent pathology. However, the contribution of constitutive antiviral defense strategies in barrier tissues, and the mechanisms by which they operate, remain poorly understood.

In this study, we report that human skin employs a pre-existing antiviral defense strategy to protect against diverse cutaneous viruses that suppress inducible IFN expression. We find that healthy human skin *in vivo* and primary skin cells cultured *in vitro* display a basal antiviral gene signature driven by the constitutive production of IFN-κ by epithelial keratinocytes.

Constitutive IFN-κ production protected keratinocytes from infection by multiple viruses, including vesicular stomatitis virus (VSV) and herpes simplex virus 1 (HSV-1), through the IFNAR. However, we also found that JAK-STAT1-dependent and -independent pathways differentially controlled replication of these viruses. Moreover, the IFN-κ-dependent, but JAK-STAT-independent, antiviral response limited HSV-1 spread to dermal fibroblasts in human skin organoids, demonstrating the ability of this cytokine to provide broad protection to multiple skin cell types. Together, our findings establish the constitutive production of IFN-κ by keratinocytes as a critical tissue autonomous antiviral defense strategy present in human skin.

## Results

### Human skin keratinocytes express an antiviral gene signature at steady state *in vivo* and *in vitro*

To determine whether human skin exhibits pre-existing antiviral defense mechanisms, we analyzed a publicly available single cell RNA-sequencing (scRNA-seq) dataset from healthy human skin(*8*). Notably, transcripts for multiple antiviral genes, including the ISGs *OAS1*, *MX1*, and *IFITM1,* were detected in the keratinocyte, fibroblast, and endothelial cell populations from these healthy biopsies (Fig. S1A). The presence of these ISG transcripts at steady state across multiple cell types in healthy human skin is consistent with constitutive production of an IFN family cytokine in this tissue. We therefore considered whether any type I or type III IFN family members were constitutively expressed in human skin. The predominant IFN cytokine detected in this tissue was *IFNK,* a type I IFN (Fig. 1A and S1B)*. IFNK* transcripts were low, but primarily detected in basal keratinocytes, consistent with reports that IFN-κ is constitutively produced by undifferentiated primary human keratinocytes *in vitro* (*9*).

**Figure 1.**
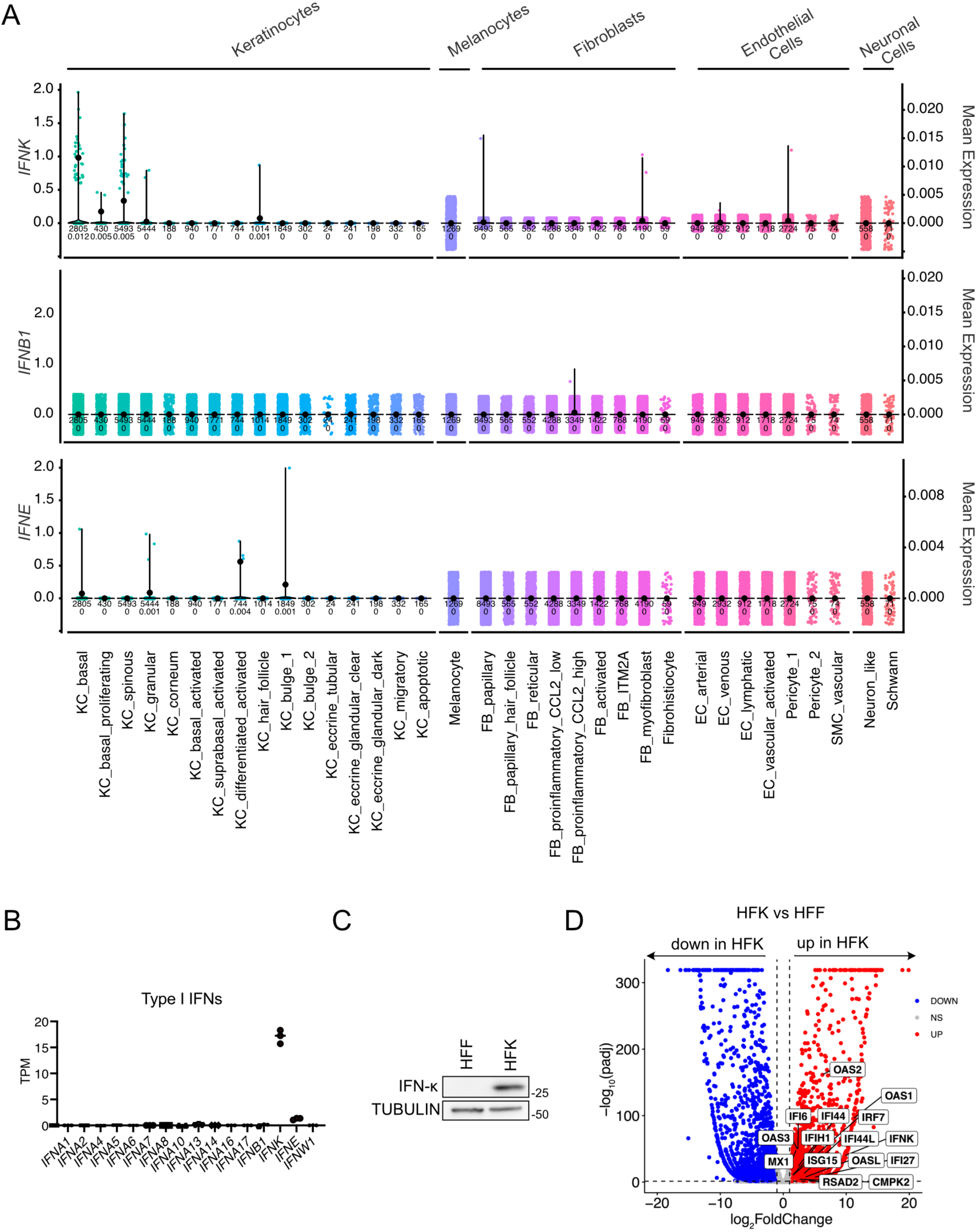
Human skin keratinocytes constitutively express *IFNK in vivo* and *in vitro*. A) Violin plots showing expression of *IFNK*, *IFNE*, and *IFNB1* in keratinocyte, melanocyte, fibroblast, endothelial cell, and neuronal cell subtypes from healthy skin punch biopsies (GSE179633). Each dot represents a single cell. B) Transcripts per million (TPM) of type-I IFN family genes quantified by bulk RNA-seq in primary human foreskin keratinocytes (HFK). C) Immunoblot of primary human foreskin fibroblasts (HFF) and HFK showing IFN-*κ* protein abundance. Images are representative of n=3 biological replicates. D) Volcano plot of differentially expressed genes in primary HFK compared to HFF. Differentially expressed genes were defined as those with a −1.0 < log_2_FC >1.0 and padj < 0.05.

To confirm that *IFNK,* and no other IFNs, are constitutively and specifically expressed by human keratinocytes *in vitro*, we performed bulk RNA-seq and immunoblot analyses on donor matched primary human keratinocyte (HFK) and fibroblast (HFF) cultures isolated from neonatal foreskin. Transcripts for *IFNK*, but not other type I IFN family members, were detected in HFK at steady state (Fig. 1B). In addition, IFN-κ protein was detectable in HFK, but not HFF, lysates (Fig. 1C). Comparison of differentially expressed genes between these two cell types, identified that *IFNK* and numerous ISGs, including *OAS1*, *OAS2*, *MX1*, *ISG15*, and *IFI44*, were among the 2,542 genes that were more highly expressed in HFK than HFF (Fig. 1D). Taken together, these findings demonstrate that an antiviral gene signature is detected in multiple cell types in healthy human skin *in vivo* and that primary keratinocytes cultured alone *in vitro* retain this phenotype.

### Constitutive *IFNK* expression in keratinocytes drives basal ISG expression and restricts diverse virus replication

The contribution of constitutive IFN-κ production to antiviral defense in human skin is poorly understood. IFN-κ has been implicated in promoting the expression of individual ISGs in keratinocytes (*9–13*) but the *IFNK*-dependent transcriptome has not been defined. To determine the contribution of IFN-κ to the global keratinocyte transcriptome and the ISG signature observed in keratinocytes *in vitro* and in human skin (Fig. 1 and S1), we performed bulk RNA-seq followed by differential gene expression analysis on RNA isolated from *IFNK*-deficient N/TERT2G keratinocytes generated by a lentiCRISPR approach. N/TERT2G cells are a normal hTERT-immortalized keratinocyte cell line that retains the barrier and biological properties of primary human keratinocytes and can be more easily manipulated by genetic engineering than primary HFK (*14*). IFN-κ protein was absent in the lysates of two N/TERT2G clones produced using distinct *IFNK*-targeting sgRNAs, sg*IFNK* #3 and #4, compared to Cas9-expressing control cells (Fig. 2A). We identified 155 downregulated genes and 44 upregulated genes in sg*IFNK* #3 N/TERT2G cells compared to control cells (Fig. S2A). Similarly, transcripts for 124 genes and 33 genes were significantly reduced or increased, respectively, in sg*IFNK* #4 N/TERT2G cells compared to control cells (Fig. S2B). There was substantial overlap between the two data sets with 106 downregulated and 16 upregulated genes shared between the two *IFNK*-deficient cell lines when compared to control cells (Fig. S2C), suggesting these are high confidence genes that are regulated by constitutive IFN-κ production.

**Figure 2.**
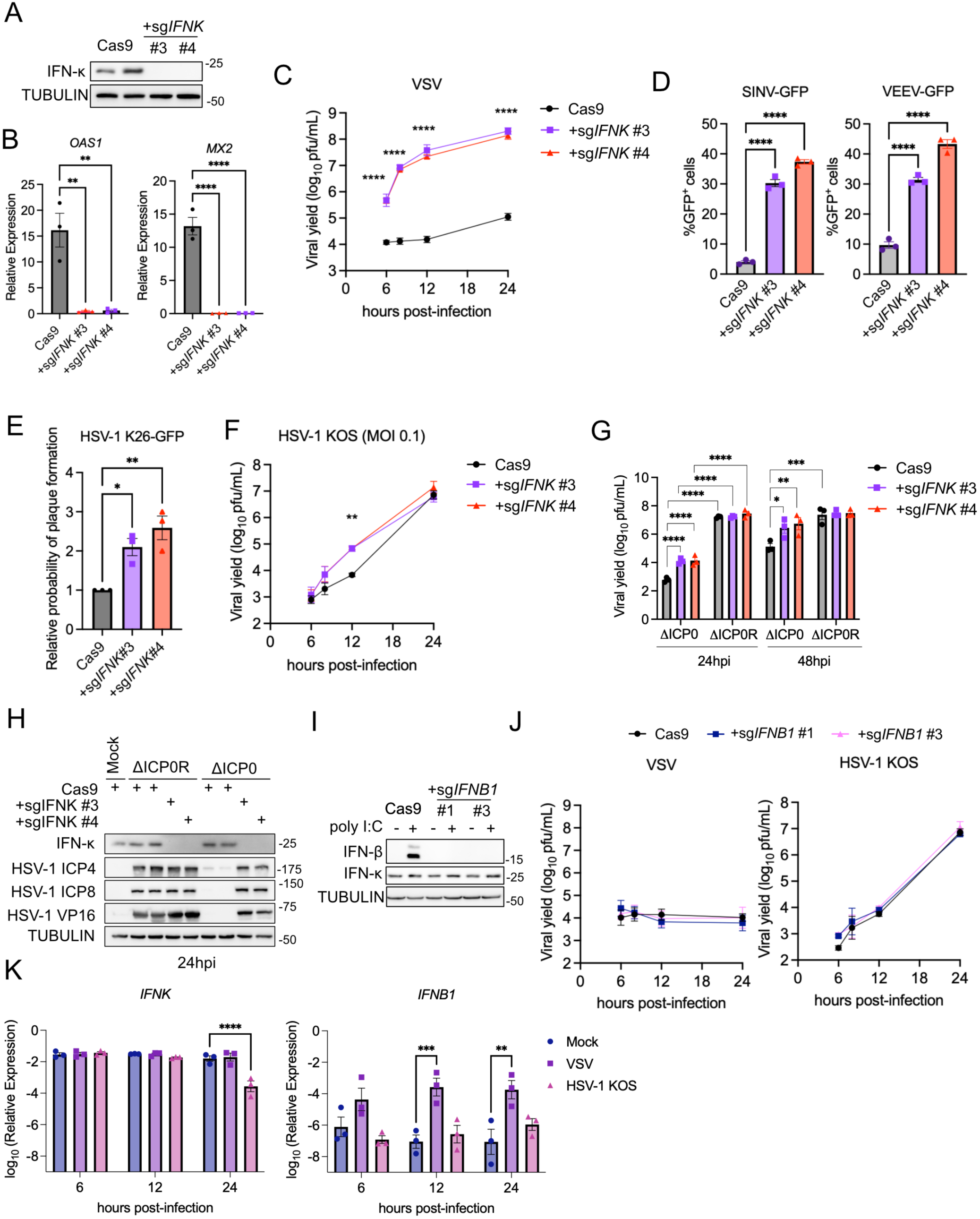
Constitutive IFNK expression in keratinocytes is broadly antiviral. A) Immunoblot of Cas9 and *IFNK*−/− N/TERT2G keratinocytes showing IFN-*κ* protein abundance. Images are representative of n=3 biological replicates. B) qRT-PCR analysis of *MX2* and *OAS1* transcript abundance in Cas9 and *IFNK*-deficient N/TERT2G keratinocytes. Data are presented as the mean ± SEM of n=3 independent experiments. C) Viral yields from Cas9 and *IFNK*-deficient N/TERT2G keratinocytes infected with VSV (MOI 1). Samples were collected for plaque assays at 6, 8, 12, and 24 h.p.i. Data are presented as the mean ± SEM of n=3 independent experiments. D) Percentage of GFP+ Cas9 and *IFNK*-deficient N/TERT2G keratinocytes at 48 h.p.i with 389-GFP or VEEV-TC-83-GFP (MOI 1). Data are presented as the mean ± SEM of n=3 independent experiments. E) Relative plaque formation in Cas9 and *IFNK*-deficient N/TERT2G keratinocytes. Data are presented as the mean ± SEM of n=3 independent experiments. F) Viral yields from Cas9 and *IFNK*−/− N/TERT2G keratinocytes infected with HSV-1 (MOI 0.1). Samples were collected for plaque assays at 6, 8, 12, and 24 h.p.i. Data are presented as the mean ± SEM of n=3 independent experiments. G) Viral yields from Cas9 and *IFNK*-deficient keratinocytes infected with HSV-1 ΔICP0 and HSV-1 ΔICP0R (MOI 0.1). Samples were collected for plaque assays at 24, and 48 h.p.i. Data are presented as the mean ± SEM of n=3 independent experiments. H) Immunoblots of lysates from Cas9 and *IFNK*-deficient N/TERT2G keratinocytes infected with HSV-1 ΔICP0 and HSV-1 ΔICP0R (MOI 10) at 24 h.p.i. I) Immunoblots of lysates from Cas9 and *IFNB1-*deficient N/TERT2G keratinocytes treated with poly I:C (10*μ*g/ml) for three hours. Images are representative of n=2 biological replicates. J) Viral yields from cas9 and *IFNB1*-deficient N/TERT2G keratinocytes infected with VSV (MOI 1) or with HSV-1 (MOI 0.1). Samples were collected for plaque assays at 6, 8, 12, and 24 h.p.i. Data are presented as the mean ± SEM of n=3 independent experiments. K) qRT-PCR analysis of *IFNK* and *IFNB1* transcript abundance relative to *18s* rRNA abundance in N/TERT2G keratinocytes infected with VSV (MOI 1) or HSV-1 (MOI 0.1) at 6,12, and 24 h.p.i. Panels C, F, G, J, K: Two-way ANOVA with Dunnett’s multiple comparisons test. Panels D and E: One-way ANOVA with Dunnett’s multiple comparisons test. *p<0.05, **p<0.01, *** p<0.001, ****p<0.0001.

We confirmed that transcripts for two *IFNK*-dependent genes, *MX2* and *OAS1*, were significantly reduced in *IFNK*-deficient keratinocytes compared to control cells by qRT-PCR (Fig. 2B). We further characterized the differentially expressed genes by gene set enrichment analysis. As expected, *IFNK*-expression was necessary for the strong type I (alpha) and type II (gamma) IFN gene signatures observed in keratinocytes (Fig. S2D-E). Unexpectedly, gene signatures associated with complement, IL6/JAK/STAT3 signaling, and TNF signaling via NFκB were also significantly reduced in both *IFNK*-deficient N/TERT2G clones (Fig. S2D-E).

Most of the *IFNK*-regulated genes identified by RNA-seq were known ISGs (*e.g*., *MX2*, *OAS1*, *RSAD2*, *IFITM1*, and *BST2*) that can be found in curated ISG databases across multiple studies (*15–18*). However, a subset of the *IFNK*-regulated genes, including *C3, LCP1, GALNT5,* and *GAMT* (Fig. S2C), are not characterized as ISGs or IFN repressed genes (IRPs). These *IFNK*-regulated genes may be induced/repressed by type I IFN stimulation specifically in keratinocytes or they might be uniquely regulated by constitutive IFN-κ production, and not by other type I IFN family members. To differentiate between these two possibilities, we examined the transcriptome of N/TERT2G cells treated with recombinant IFN-β for 6 hrs by bulk RNA-seq. We identified 345 upregulated and 46 downregulated genes in IFN-β stimulated N/TERT2G keratinocytes (Fig. S2F). When we compared the IFN-β and IFN-κ regulated transcriptomes, we found substantial overlap between the datasets, including the classic ISGs *MX2* and *OAS1* (Fig S2G-I). However, we also identified a subset of IFN-κ-regulated genes (42/122 genes) that were not similarly regulated by IFN-β treatment (Fig. S2G). This included the IFN-κ-stimulated genes *C3* and *LCP1*, and all 16 of the IFN-κ-repressed genes (Fig. S2H and I). Thus, constitutive IFN-κ production in keratinocytes appears to drive a unique transcriptional profile in these cells that is not completely shared by recombinant IFN-β treatment.

Since ISG expression correlates with resistance to virus infection (*18–21*) we next tested if virus replication was enhanced in *IFNK-*deficient N/TERT2G cells compared to control cells. We tested four RNA and DNA viruses from diverse families that use skin as a portal for entry and can infect human keratinocytes, including vesicular stomatitis virus (VSV)(*22, 23*), herpes simplex virus 1 (HSV-1)(*24*), Sindbis virus (SINV)(*25*), and Venezuelan equine encephalitis virus (VEEV)(*26*). VSV replication was potently controlled by the presence of IFN-κ in N/TERT2G cells, as VSV titers were enhanced by 2-4 logs at multiple time points in *IFNK*-deficient cells, when compared to control cells (Fig. 2C). The percentages of N/TERT2G cells infected with GFP-expressing SINV or VEEV reporter viruses were also significantly increased in *IFNK-*deficient cells when compared to control cells (Fig. 2D), suggesting that *IFNK* can control the replication of multiple RNA viruses that infect keratinocytes.

The DNA virus HSV-1 was more capable of establishing infection in keratinocytes in the absence of IFN-κ, as demonstrated by the increased probability of a GFP-reporter virus (K26-GFP) to form plaques on *IFNK*-deficient N/TERT2G monolayers than those generated with control cells (Fig. 2E). Further, when we assessed multi-cycle replication kinetics of the HSV-1 KOS strain in these cells over time, we found that viral titers were transiently enhanced (∼1 log) at 12 hpi in *IFNK*-deficient cells compared to control cells (Fig. 2F). The transient nature of this restriction may reflect the numerous innate immune evasion strategies employed by this virus that facilitate its replication in the presence of IFNs (*5, 27–34*). We therefore also examined the contribution of *IFNK* to the restriction of an IFN-sensitive HSV-1 mutant lacking the viral protein ICP0 (HSV-1 ΔICP0)(*35*), which we have previously shown has a profound replication defect in primary human keratinocytes at low multiplicity of infection (MOI) (*36*). Like our observations in primary cells, HSV-1 ΔICP0 replicates poorly in N/TERT2G cells compared to a revertant virus where ICP0 expression has been restored (HSV-1 ΔICP0R) (Fig. 2G). The replication defect of HSV-1 ΔICP0 was partially restored in *IFNK*-deficient N/TERT2G cells at 24 hours post-infection (hpi) and fully restored at 48 hpi (Fig. 2G). When compared to HSV-1 ΔICP0R infection at high MOI (10 pfu/cell), which allows for a single cycle of virus replication to be examined, HSV-1 ΔICP0 infection of control cells resulted in limited or no detection of viral proteins ICP4, ICP8, and VP16 at 24 hpi (Fig. 2H). By contrast, all three proteins were substantially increased in HSV-1 ΔICP0-infected *IFNK*-deficient cell lysates compared to Cas9-control cells (Fig. 2H). The abundance of these proteins in HSV-1 ΔICP0 infected *IFNK*-deficient cell lysates was equivalent to that observed in revertant virus infected cells, suggesting that IFN-κ limits HSV-1 ΔICP0 replication in human keratinocytes by reducing HSV-1 gene expression.

To determine if enhanced replication of these diverse viruses was specific to *IFNK* expression, we next assessed multi-cycle replication kinetics of VSV and HSV-1 in *IFNB1*-deficient N/TERT2G cells. Because *IFNB1* is not expressed at steady state in keratinocytes (Fig. 1B), we validated CRISPR-mediated *IFNB1*-deficiency following treatment with poly I:C, a TLR3 agonist that induces *IFNB1* expression (*37, 38*). Poly I:C treatment induced the accumulation of IFN-β protein in control, but not *IFNB1*-deficient N/TERT2G cells, and had no effect on IFN-κ abundance (Fig. 2I). We observed no increase in VSV or HSV-1 yields in *IFNB1*-deficient N/TERT2G cells at any time point examined (Fig. 2J), consistent with these viruses limiting inducible type I IFN production/function in keratinocytes (Fig. 2K)(*23, 39*). These data together suggest that constitutive IFN-κ production in keratinocytes drives a basal antiviral transcriptional signature and restricts the replication of diverse cutaneous viruses.

### A canonical IFNAR-dependent pathway controls VSV, but not HSV-1, replication in human keratinocytes

Recombinant IFN-κ drives an IFNAR1/2-dependent pathway in A549 lung epithelial cells that reduces influenza virus replication (*40*), so we investigated the type I IFN signaling pathway components that are important for constitutive *IFNK*-dependent control of VSV and HSV-1 replication in keratinocytes. We first examined ISG expression and virus replication in *IFNAR1-* and *IFNAR2-*deficient N/FTERT2G keratinocytes (Fig. S3A). Expression of the *IFNK*-dependent gene *MX2* (Fig. 2B, S2H) was significantly reduced in multiple clones of *IFNAR1-* and *IFNAR2*-deficient N/TERT2G keratinocytes in the absence of infection (Fig. 3A). In addition, like our observations in *IFNK*-deficient N/TERT2G cells (Fig. 2), the absence of *IFNAR1* or *IFNAR2* in N/TERT2G cells significantly increased VSV (∼4 orders of magnitude) and HSV-1 KOS (∼1 order of magnitude) yields when compared to Cas9-control cells (Fig. 3B and C). Further, HSV-1 ΔICP0 virus replication was partially restored in *IFNAR1-* and *IFNAR2*-deficient N/TERT2G cells at 24 and 48 hpi (Fig. 3D). Together these observations suggest that constitutive IFN-κ-mediated restriction of VSV and HSV-1 in keratinocytes requires both the IFNAR1 and IFNAR2 subunits.

**Figure 3.**
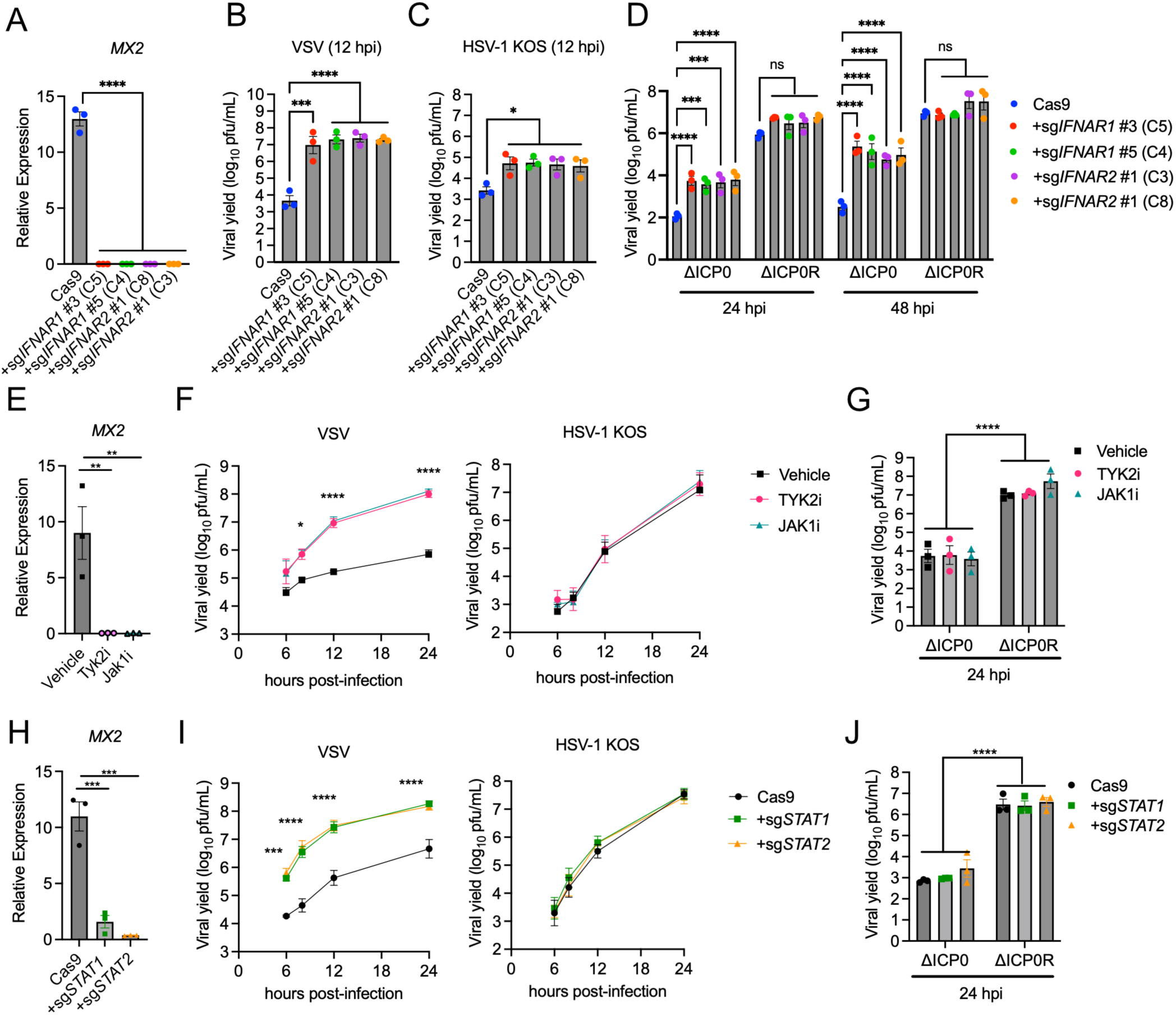
Tonic JAK-STAT signaling differentially controls VSV and HSV-1 replication in human keratinocytes. A) qRT-PCR analysis of *MX2* transcript abundance relative to *TBP* in control (Cas9), *IFNAR1*-, and *IFNAR2*-deficient N/TERT2G keratinocytes. Data are presented as the mean ± SEM of n=3 independent experiments. B and C) Viral yields from Cas9, *IFNAR1*-deficient, and *IFNAR2*-deficient N/TERT2G keratinocytes infected with B) VSV (MOI 1.0) or C) HSV-1 KOS (MOI 0.1). Samples were collected at 12 hpi. Data are presented as the mean ± SEM of n=3 independent experiments. D) Viral yields from Cas9, *IFNAR1*-deficient, and *IFNAR2*-deficient N/TERT2G keratinocytes infected with HSV-1 ΔICP0 and HSV-1 ΔICP0R (MOI 0.1). Samples were collected at 24 and 48 hpi. Data are presented as the mean ± SEM of n=3 independent experiments. E) qRT-PCR analysis of *MX2* transcript abundance relative to *TBP* in N/TERT2G keratinocytes treated with 10 *μ*M of Abrocitinib (JAK1i) or Deucravacitinib (TYK2i) for 24 hrs. Data are presented as the mean ± SEM of n=3 independent experiments. F and G) Viral yields from N/TERT2G keratinocytes treated with 10 *μ*M of Abrocitinib (JAK1i) or Deucravacitinib (TYK2i) 24 hours prior to infection with F) VSV (MOI 1) or HSV-1 KOS (MOI 0.1) or G) HSV-1 ΔICP0 and HSV-1 ΔICP0R (MOI 0.1). Samples were collected at 6, 8,12, and 24 hpi for HSV-1 and VSV. Samples were collected at 24 hpi for HSV-1 ΔICP0 and HSV-1 ΔICP0R. Data are presented as the mean ± SEM of n=3 independent experiments. H) qRT-PCR analysis of *MX2* transcript abundance relative to *TBP* in Cas9, *STAT1-*, and *STAT2-*deficient N/TERT2G keratinocytes. Data are presented as the mean ± SEM of n=3 independent experiments. I) Viral yields from Cas9, *STAT1-*deficient and *STAT2-*deficient N/TERT2G keratinocytes infected with VSV (MOI 1) or HSV-1 KOS (MOI 0.1). Samples were collected at 6, 8,12, and 24 hpi. Data are presented as the mean ± SEM of n=3 independent experiments. J) Viral yields from Cas9, *STAT1-*deficient, and *STAT2-*deficient N/TERT2G keratinocytes infected with HSV-1 ΔICP0 and HSV-1 ΔICP0R (MOI 0.1) for 24 hrs. Data are presented as the mean ± SEM of n=3 independent experiments. Panels A-C, E, H: One-way ANOVA with Tukey’s multiple comparisons test. Panels D, F-G, and I-J: Two-way ANOVA with Dunnett’s multiple comparisons test. *p<0.05, **p<0.01, *** p<0.001, ****p<0.0001.

The canonical type I IFN signaling pathway requires the kinase function of JAK family members TYK2 and JAK1, which phosphorylate STAT1 and STAT2 to promote their activation and drive ISG expression. We therefore investigated the requirement for JAK-STAT signaling in tonic ISG expression and virus restriction in keratinocytes. Treatment with TYK2 and JAK1 kinase inhibitors abrocitinib (JAK1i) and deucravacitinib (TYK2i) individually reduced the steady state expression of *MX2* (Fig. 3E) and increased VSV yields by multiple orders of magnitude (Fig. 3F). By contrast, neither WT HSV-1 (KOS or ICP0R) nor ΔICP0 virus replication were enhanced in TYK2 or JAK1 inhibitor treated keratinocytes (Fig. 3F and G), suggesting that constitutive *IFNK*-dependent control of VSV and HSV-1 replication in keratinocytes rely on distinct signaling components (*41*).

To determine the contribution of STAT1 and STAT2 to constitutive antiviral defense in keratinocytes we generated *STAT1*- or *STAT2*-deficient N/TERT2G keratinocytes using CRISPR. By immunoblot analysis, lysates from a pooled population of *STAT1*-deficient cells showed reduced STAT1 abundance with limited effects on STAT2 or IFN-κ (Fig. S3B). However, deletion of *STAT2* in N/TERT2G cells resulted in a reduction in both STAT2 and STAT1 protein abundance. This observation suggests that steady-state expression of *STAT2* in keratinocytes drives the expression or stability of STAT1 and is consistent with our observations that both *STAT1* and *STAT2* expression in keratinocytes are enhanced by constitutive IFN-κ signaling (Fig. S2C). Similar to our observations with kinase inhibitor treatment, *STAT1-* and *STAT2*-deficient N/TERT2G cells showed reduced *MX2* expression (Fig. 3H) and increased VSV yields at all time points examined (Fig. 3I). However, the absence of either of these two transcription factors had no effect on WT HSV-1 (KOS or ΔICP0R) or HSV-1 ΔICP0 yields (Fig. 2I and J).

These data together suggest that distinct IFNAR-dependent signaling pathways control VSV and HSV-1 replication in keratinocytes. VSV infection is controlled by a canonical JAK-STAT-dependent pathway in keratinocytes, while keratinocytes use an IFNAR1-dependent, but JAK-STAT-independent mechanism to limit HSV-1 replication.

### Keratinocyte-derived IFN-κ protects primary human fibroblasts from HSV-1 infection in a skin organoid model

In monoculture systems *in vitro*, primary human keratinocytes and fibroblasts isolated from human skin are generally readily infected with HSV-1 (Fig. 2F and J) (*23, 27, 36*). However, dermal fibroblasts in skin explants exposed to HSV-1 exhibit limited infection (*42–44*). These published findings, together with our observations that human keratinocytes protect themselves from virus infection via constitutive IFN-κ production (Fig. 2) and that fibroblasts *in vivo* exhibit a basal ISG signature (Fig. S1), led us to hypothesize that keratinocyte-derived IFN-κ may also protect dermal fibroblasts from virus infection in human skin. To test this possibility, we sought to establish an *in vitro* skin-like system that recapitulated the basal ISG signature observed in human skin fibroblasts *in vivo*. We generated two organotypic human skin models: human skin equivalents (HSE) and dermal equivalents (DE). HSE are organoids composed of primary human keratinocytes at various stages of differentiation, grown at an air-liquid interface atop a collagen matrix containing primary human fibroblasts (Fig. 4A and B) (*45, 46*). By contrast, DE organoids are built similarly to HSE organoids but without the addition of keratinocytes. Parallel generation of DE and HSE organoids followed by targeted qRT-PCR analysis of fibroblasts isolated from these cultures, revealed increased expression of multiple ISGs, including *MX2* and *BST2*, in HSE fibroblasts relative to DE fibroblasts (Fig. 4C). These findings indicated that HSE organoids recapitulate the basal ISG signature observed in human skin fibroblasts *in vivo* and suggest that keratinocytes are sufficient to drive this response during organoid development.

**Figure 4.**
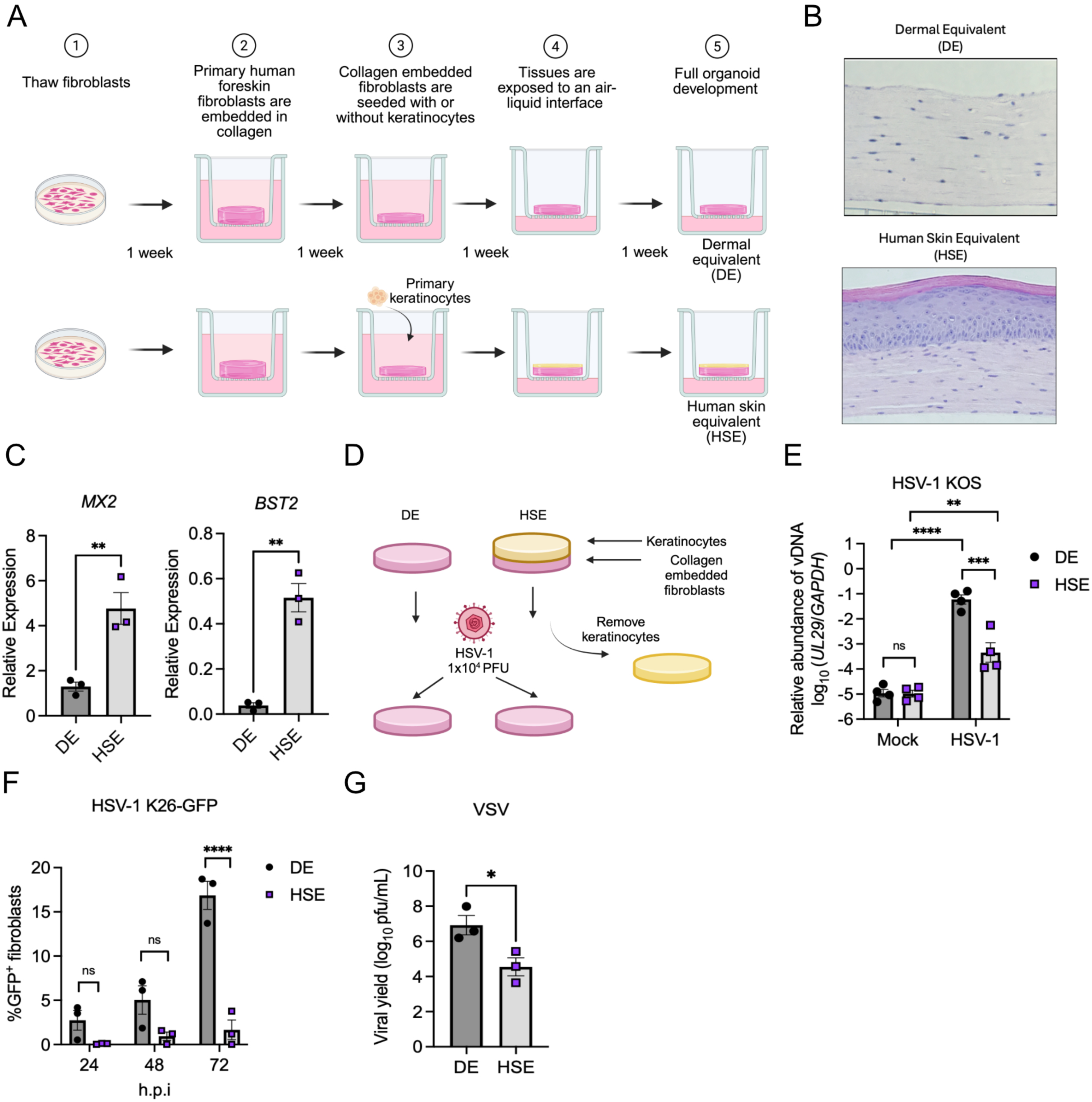
Fibroblasts cultured with keratinocytes in human skin organoids are resistant to virus infection. A) Overview of skin organoid construction. (Top) Dermal equivalent (DE) organoids consist of fibroblasts embedded in collagen, forming a dermis-like tissue. (Bottom) Human skin equivalent (HSE) organoids contain keratinocytes seeded on top of dermal fibroblasts embedded in collagen. The tissues are exposed to an air-liquid interface to promote epithelial differentiation. Created in BioRender. Johnson, J. (2026) https://BioRender.com/nxjs6n8. B) Hematoxylin and eosin staining of HSE and DE cross sections showing the morphological characteristics of the skin organoids. Scale bar = 570 μm. C) qRT-PCR analysis of *MX2* and *BST2* transcript abundance relative to *TBP* in DE and HSE fibroblasts Data are presented as the mean ± SEM of n=3 independent experiments. D) Schematic demonstrating the experimental approach used to study basal antiviral communication between keratinocytes and fibroblasts. Created in BioRender. Johnson, J. (2026) https://BioRender.com/nbw6srz. E) qPCR analysis of HSV-1 KOS viral DNA (vDNA) (*UL29*) relative to cellular DNA *(GAPDH)* in DE or HSE fibroblasts at 24 hpi. DE and HSE fibroblasts were infected with 1×10^4^ pfu of HSV-1 KOS. Data are presented as the mean ± SEM of n=4 independent experiments. F) Percent of GFP+ DE or HSE fibroblasts at 24, 48, and 72 hpi after infection with HSV-1 K26-GFP (1×10^4^ pfu). G) Viral yields from supernatants of DE and HSE fibroblasts infected with VSV (1×10^4^ pfu) at 24 hpi. Panels E and F: Two-way ANOVA with Sidak’s multiple comparisons test. Panels C and G: Unpaired t test. *p<0.05, **p<0.01, *** p<0.001, ****p<0.0001.

To determine whether keratinocytes confer protection to fibroblasts against HSV-1 in our organoid models, we first generated DE and HSE cultures in parallel. After the development of a fully differentiated epidermis in HSE cultures, as demonstrated by the formation of cornified skin (Fig. 4B), we enzymatically removed the epidermis and directly infected the collagen embedded dermal fibroblasts from HSE and DE cultures with HSV-1 (Fig. 4D). To assess infection, we quantified viral DNA abundance (vDNA) in organoid fibroblasts at 24 hpi by qPCR for the HSV-1 gene *UL29*. The relative abundance of HSV-1 vDNA was ∼ 2.5 orders of magnitude more abundant in DE fibroblasts than in HSE fibroblasts (Fig. 4E). To determine the proportion of infected fibroblasts, we infected DE and HSE fibroblasts with the HSV-1 K26-GFP reporter virus (*47*) and quantified GFP positive fibroblasts by flow cytometry (Fig. S4). Consistent with vDNA measurements, a higher percentage of GFP positive fibroblasts was observed in DE cultures than in HSE cultures at all time points examined, reaching statistical significance at 72 hpi (Fig. 4F). Restriction of virus replication in HSE fibroblasts compared to DE fibroblasts was not limited to HSV-1, as we also observed reduced VSV viral yields in HSE fibroblasts (Fig. 4G). Collectively, these experiments establish an *in vitro* organoid system that recapitulates the relative resistance of dermal fibroblasts to HSV-1 infection observed in human skin explants (*42–44*). Further, these data suggest that the presence of keratinocytes prior to infection reduces the ability of diverse viruses to efficiently infect dermal fibroblasts.

The presence of a keratinocyte-dependent ISG signature and restriction of HSV-1 replication in HSE fibroblasts prompted us to determine if constitutive IFN-κ production by keratinocytes underlies these phenotypes. To test this, we generated HSE organoids using control (WT) or *IFNK^−/−^* N/TERT2G keratinocytes (Fig. 5A), in parallel with DE cultures, and quantified steady-state ISG transcript abundance and HSV-1 replication in organoid fibroblasts (Fig. 5B). HSEs constructed with *IFNK^−/−^* N/TERT2G keratinocytes exhibited morphological characteristics comparable to those generated with WT cells (Fig. 5C), indicating that keratinocyte-derived IFN-κ does not overtly affect skin organoid development or differentiation. Consistent with our findings in HSEs generated with primary keratinocyte cultures (Fig. 4), fibroblasts in HSE organoids constructed with WT N/TERT2G keratinocytes exhibited increased expression of *BST2* and *MX2* in the absence of infection (Fig. 5D) and reduced HSV-1 vDNA abundance following infection (Fig. 5E) relative to DE fibroblasts. By contrast, both the elevated ISG signature and the restriction of HSV-1 in HSE fibroblasts generated with *IFNK^−/−^* N/TERT2G keratinocytes were comparable to that seen in DE fibroblasts (Fig. 5D and E). Together, these results demonstrate that constitutive production of IFN-κ by epidermal keratinocytes promotes basal ISG expression and establishes an antiviral state in neighboring dermal fibroblasts.

**Figure 5.**
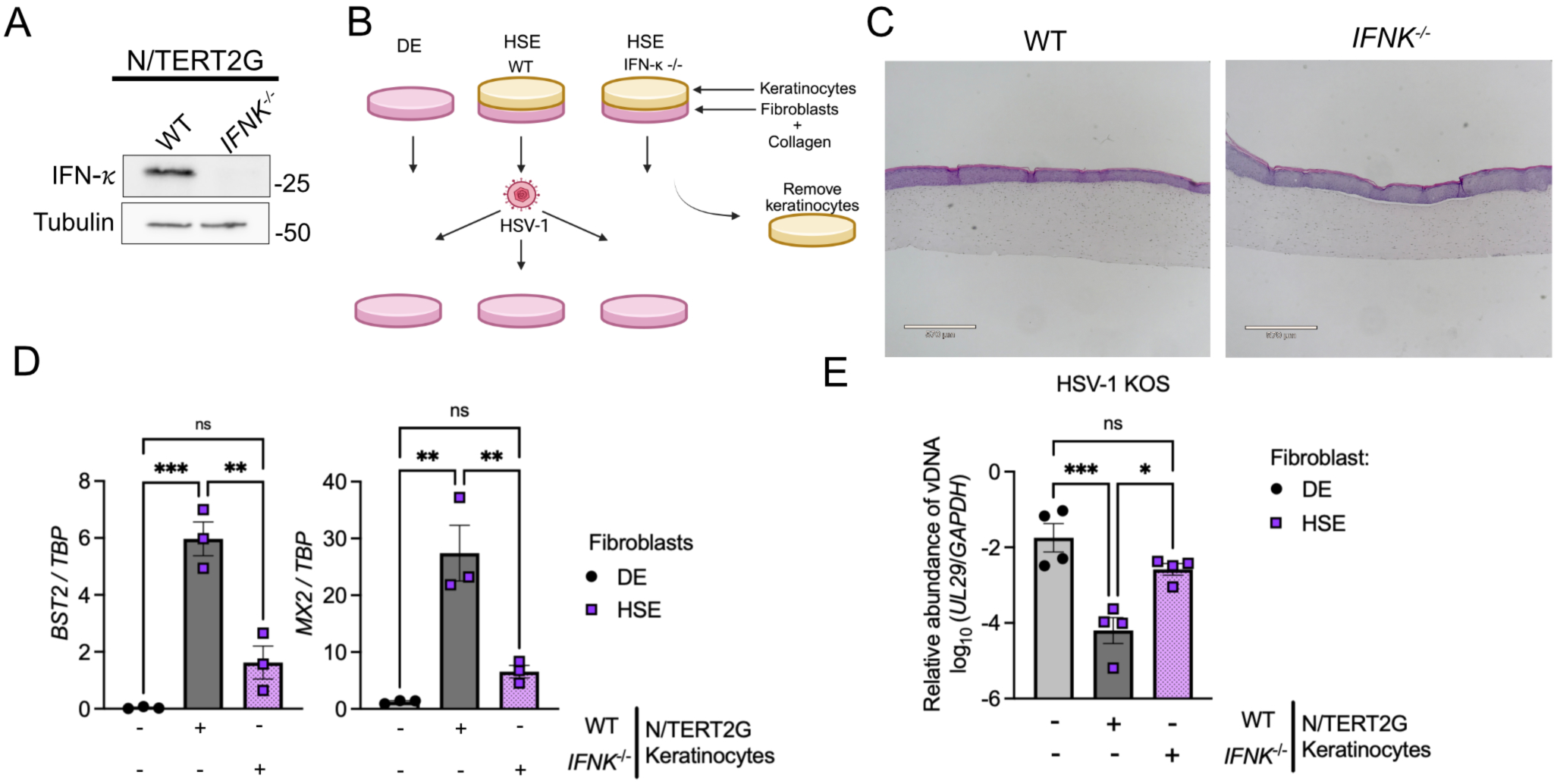
Keratinocyte-derived IFN-k drives an antiviral response in skin organoid fibroblasts. A) Immunoblots of lysates from wild-type (WT) and *IFNK^−/−^*N/TERT2G keratinocytes showing IFN-*κ* protein abundance. Images are representative of n=3 biological replicates. B) Depiction of experimental design. (Top) DE and HSE organoids generated with WT or *IFNK*^−/−^ N/TERT2G keratinocytes were constructed in parallel. HSE keratinocytes were removed and fibroblasts from all organoid cultures were infected with 1×10^4^ pfu of HSV-1 KOS. Created in BioRender. Johnson, J. (2026) https://BioRender.com/7s24bz0. C) Hematoxylin and eosin staining of HSE cross sections from WT and *IFNK*^−/−^ organoid cultures showing the morphological characteristics of the organoids. Scale bar = 570 μM. D) *MX2* and *BST2* transcript abundance relative to *TBP* in DE, WT HSE, and *IFNK*^−/−^ HSE organoid fibroblasts. Data are presented as the mean ± SEM of n=3 independent experiments. E) qPCR analysis of HSV-1 KOS vDNA (*UL29*) abundance relative to cellular DNA *(GAPDH)* in DE, WT HSE, and *IFNK*^−/−^ HSE organoid fibroblasts at 24 hpi. DE and HSE fibroblasts were infected with 1×10^4^ pfu/ml of HSV-1 KOS. Data are presented as the mean ± SEM of n=4 independent experiments. Panels D and E: One-Way ANOVA with Tukey’s multiple comparison test. *p<0.05, **p<0.01, *** p<0.001, ****p<0.0001.

### Keratinocyte-mediated restriction of HSV-1 in fibroblasts is not dependent on a canonical *IFNAR*-dependent pathway

Given that HSV-1 restriction in HSE fibroblasts is dependent on keratinocyte-derived IFN-κ and that this cytokine restricts HSV-1 replication in keratinocytes via a non-canonical mechanism (Fig. 3 and 4), we next asked whether IFN-κ similarly drives a non-canonical antiviral pathway in HSE fibroblasts. To test the contribution of IFNAR and canonical signaling components to antiviral defense in HSE organoids, we used CRISPR to generate hTERT-fibroblasts deficient in *IFNAR1*, *STAT1*, *STAT2* or *IRF9* (Fig. 6A) and incorporated these cells in DE and HSE organoids for analysis. Before organoid generation, we confirmed impaired type I IFN signaling in these CRISPR-modified hTERT-fibroblast lines by treating them with recombinant IFNs and assessing STAT1 phosphorylation in the *IFNAR1*-deficient cells or restriction of the IFN-sensitive virus VSV in *STAT1-*, *STAT2-,* or *IRF9*-deficient cells. In Cas9-control hTERT-fibroblasts, both IFN-β and IFN-ɣ, a type II IFN that does not utilize IFNAR for signaling, induced STAT1 phosphorylation (Fig. S5A and B). By contrast, *IFNAR1*-deficient fibroblasts generated with two distinct sgRNAs failed to induce robust STAT1 phosphorylation following IFN-β treatment while retaining their responsiveness to IFN-ɣ (Fig. S5A and B). In addition, IFN-β suppressed VSV replication in Cas9-control hTERT-fibroblasts, but not in *STAT1-*, *STAT2*-, or *IRF9*-deficient cells (Fig. S5C).

**Figure 6.**
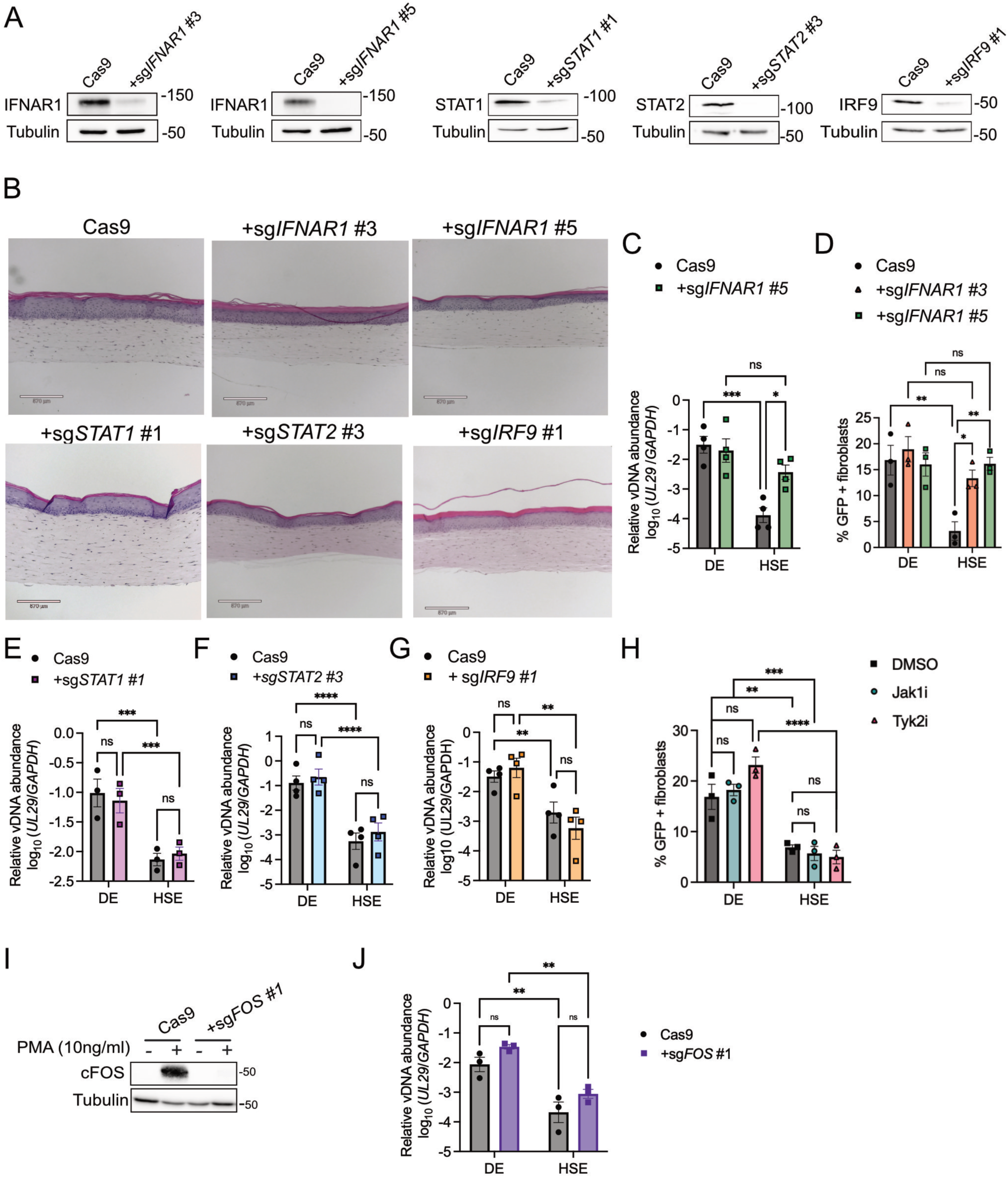
Restriction of HSV-1 in HSE fibroblasts is independent of canonical IFNAR signaling components. A) Immunoblots of lysates from control (Cas9), *IFNAR1*-, *STAT1*-, *STAT2*-, and *IRF9*-deficient fibroblasts demonstrating knockout efficiency of target genes. B) Hematoxylin and eosin staining of HSE sections showing the morphological characteristics of organoid cultures generated with Cas9, *IFNAR1*-, *STAT1*-, *IRF9*-, and *STAT2*-deficient fibroblasts. Scale bar = 570 μM. C) qPCR analysis of HSV-1 KOS vDNA (*UL29*) abundance relative to cellular DNA (*GAPDH*) in Cas9 and *IFNAR1*-deficient organoid fibroblasts infected with 1×10^4^ pfu of HSV-1 KOS for 24 hrs. D) Flow cytometry analysis quantifying the percent of GFP positive Cas9 or *IFNAR1*-deficient organoid fibroblasts infected with 1×10^4^ pfu HSV-1 K26-GFP at 72 hpi. Data are presented as the mean ± SEM of n ≥ 3 independent experiments. E-G) qPCR analysis of HSV-1 KOS vDNA (*UL29*) abundance relative to cellular DNA (*GAPDH*) abundance in Cas9, *STAT1*-, *STAT2*-, and *IRF9*-deficient fibroblasts cultured in DE and HSE organoids. Fibroblasts were infected with 1×10^4^ pfu of HSV-1 KOS for 24 hrs. Data are presented as the mean ± SEM of n=3 independent experiments. H) Percent of GFP positive DE or HSE fibroblast at 72 hpi after infection with HSV-1 K26-GFP (1×10^4^ pfu). Organoid cultures were treated with 10 *μ*M of Abrocitinib (Jaki) or Deucravacitinib (Tyk2i) for 24 hours prior to infection. Data are presented as the mean ± SEM of n=3 independent experiments. I) Immunoblots of lysates from control (Cas9) and *FOS*-deficient fibroblasts treated with PMA (10 ng/ml) for 30 minutes demonstrating knockout efficiency. J) qPCR analysis of HSV-1 KOS vDNA (*UL29*) abundance relative to cellular DNA (*GAPDH*) abundance in Cas9 and *FOS*-deficient fibroblasts cultured in DE and HSE organoids. Fibroblasts were infected with 1×10^4^ pfu of HSV-1 KOS for 24 hrs. Data are presented as the mean ± SEM of n=3 independent experiments.

HSE organoids generated with *IFNAR1-*, *STAT1-*, *STAT2-,* or *IRF9*-deficient hTERT-fibroblasts exhibited morphology comparable to that of organoids generated with Cas9-control cells (Fig. 6B). To determine how loss of IFNAR signaling affected the enhanced *IFNK*-dependent ISG signature we observed in HSE fibroblasts relative to DE fibroblasts (Fig. 4C and 5D), we first performed bulk RNA-seq followed by differential gene expression analysis on Cas9-control hTERT-fibroblasts and one *IFNAR1*-deficient hTERT-fibroblast line isolated from uninfected HSE cultures. We identified 25 genes that were significantly down regulated in *IFNAR1*-deficient HSE fibroblasts relative to control HSE fibroblasts, including canonical ISGs *MX2*, *IFITM1,* and *OAS3* (Fig. S5D). Notably, the non-canonical *IFNK*-dependent ISGs and IRGs that we identified in keratinocytes (Fig. S2) were not differentially expressed in *IFNAR1*-deficient HSE fibroblasts, possibly reflecting the inability of bulk RNAseq analysis to capture changes in gene expression that likely occur in a spatially defined manner in these tissues.

To define the signaling components required for the canonical ISG response we observed in HSE fibroblasts, we next performed targeted RT-qPCR analysis of organoids generated with *STAT1-*, *STAT2-*, or *IRF9*-deficient fibroblasts. These experiments revealed that *STAT2* and *IRF9* were essential for maintaining keratinocyte induced expression of the canonical ISGs *MX2, OAS3,* and *BST2* in uninfected HSE fibroblasts (Fig. S5E-F). By contrast, *STAT1* deficiency had a minor effect on *MX2* and *OAS3*, but not *BST2,* expression (Fig. S5G). Previous studies in other cell types have shown that STAT2:IRF9 heterodimers can drive constitutive and transient ISG responses (*48–50*). Thus, STAT2:IRF9 heterodimers may be the predominant contributors to the keratinocyte-dependent canonical ISG response in HSE fibroblasts.

We next compared the ability of HSV-1 KOS or HSV-1 K26-GFP to infect Cas9-control or *IFNAR1-*, *STAT1-*, *STAT2-* and *IRF9*-deficient fibroblasts in DE and HSE cultures. Consistent with our prior observations using unmodified primary fibroblasts (Fig. 3), HSV-1 KOS vDNA abundance was reduced by ∼2.5 orders of magnitude in Cas9-control hTERT-fibroblasts from HSE cultures relative to DE cultures (Fig. 6D). We likewise observed fewer HSV-1 K26-GFP-infected fibroblasts in HSE cultures generated with Cas9-control hTERT-fibroblasts than in DE cultures (Fig. 6E). By contrast, both HSV-1 KOS vDNA abundance and the percentage of HSV-1 K26-GFP infected fibroblasts were significantly increased in *IFNAR1*-deficient HSE fibroblasts, reaching levels comparable to those observed in DE fibroblasts (Fig. 6D and E). Notably, *IFNAR1*-deficiency had no effect on vDNA abundance or GFP + fibroblast numbers in DE cultures, consistent with the inability of human fibroblasts to mount a functional antiviral type I IFN response to HSV-1 infection in the absence of keratinocytes (*27, 34, 51, 52*). Despite the requirement for *IFNAR1* expression in restricting HSV-1 replication in HSE fibroblasts, loss of *STAT1*, *STAT2*, or *IRF9* had no effect on HSV-1 replication in HSE or DE fibroblasts (Fig. 6F-H). Similarly, although treatment of organoid cultures with JAK1 or TYK2 inhibitors reduced keratinocyte-induced *MX2* expression in fibroblasts (Fig. S5H), these treatments did not alter HSV-1 infection of fibroblasts in organoids (Fig. 5I).

IFN-κ can control IAV replication in lung epithelial cells through an IFNAR-dependent, but STAT1-independent, pathway that requires the transcription factor c-Fos (*40*), which in turn drives expression of *CHD6*, the principal effector of this cytokine’s anti-IAV activity. *CHD6* was not among the IFNAR1-dependent genes identified in HSE fibroblasts; however, *FOS*, which encodes c-Fos, was significantly reduced in *IFNAR1*-deficient HSE fibroblasts relative to control cells (Fig. S5D). We therefore asked whether *FOS* expression in fibroblasts was required for IFN-κ/IFNAR1-dependent inhibition of HSV-1 replication in organoid fibroblasts. To this end, we generated *FOS*-deficient hTERT fibroblasts by CRISPR-Cas9. c-Fos protein was undetectable at steady-state in fibroblast monocultures but was inducible by treatment with phorbol 12-mytrstate 13-acetae (PMA), confirming loss of c-Fos protein in *FOS*-deficient cells (Fig. 6H). HSV-1 vDNA abundance was unaffected by loss of c-Fos, indicating that changes in *FOS* expression could not explain the IFNAR1-dependent control of HSV-1 (Fig. 6I). Together, these results suggest that, as in keratinocytes, IFN-κ/IFNAR1-dependent control of HSV-1 in organoid fibroblasts operates through a mechanism that is distinct from both canonical JAK/STAT-dependent signaling and the previously described non-canonical c-Fos-dependent program of ISG induction.

## Discussion

In this study, we demonstrated that constitutive IFN-κ production by human keratinocytes establishes a preexisting antiviral state in human skin (Fig. S6). Constitutive expression of an IFN family member in the epidermis of normal human skin was proposed several decades ago (*53*) and was later identified as IFN-κ (*9*). Although IFN-κ has been implicated in autoinflammatory skin disease(*13, 54*), its role in cutaneous antiviral defense has remained unclear. Early studies showed that recombinant IFN-κ restricts encephalomyocarditis virus replication in fibroblasts and induces ISG expression in a human b-lymphoblastoid cell line (*9*). Subsequent work using doxycycline-inducible expression systems showed that IFN-κ reduces human papillomavirus transcription and replication (*11*), suggesting that supraphysiological levels of IFN-κ can be antiviral. Here, we extend these findings by showing that constitutive IFN-κ production is a functional component of the innate immune system in human skin and profoundly limits the replication of diverse viruses, including VSV, HSV-1, SINV, and VEEV in human keratinocytes. Moreover, this constitutive antiviral activity is not restricted to keratinocytes, but extends to proximal cell types in human skin, as dermal fibroblasts in skin organoids are protected from HSV-1 infection in an epidermal IFN-κ-dependent manner.

Our investigation of the signaling pathways downstream of constitutive IFN-κ production identified notable virus-specific differences in the requirement for canonical signaling components in viral restriction. We found that VSV restriction depended on keratinocyte expression of the type-I IFN receptor subunits IFNAR1/2, the activity of the receptor proximal kinases JAK1 and TYK2, and the downstream transcription factors STAT1 and STAT2. This comprehensive analysis of the IFNAR signaling pathway suggests that constitutive IFN-κ-mediated restriction of VSV follows the canonical JAK-STAT signaling pathway, consistent with the reported anti-VSV activity of recombinant IFN-β (*55–58*). However, these same canonical components were not required for IFN-κ-mediated restriction of HSV-1 in keratinocytes, suggesting the existence of a non-canonical antiviral pathway downstream of this cytokine. Moreover, the IFN-κ−/IFNAR1-mediated HSV-1 restriction in dermal fibroblasts was also JAK-STAT independent, arguing that the putative non-canonical signaling pathway engaged by IFN-κ is not cell type specific, but rather a feature of IFN-κ signaling in human skin.

A protective role for a JAK-STAT-independent pathway in HSV-1 skin infection is notable for several reasons. First, this virus encodes several proteins that target canonical PRR-induced IFN signaling pathway components to evade the host innate immune response(*5, 27–34*). Thus, a pathway that does not rely on canonical signaling components may serve as a mechanism by which the innate immune system limits the negative impact of HSV-1 immune evasion strategies. Second, JAK-STAT-independent control of HSV-1 in human skin may explain why patients treated with oral JAK inhibitors show no increased risk of HSV infections but are susceptible to other virus infections (44). More broadly, our findings may have therapeutic implications for the use of immunomodulators that target JAK family members in the context of cutaneous viral infections.

The IFN-κ-dependent, but JAK-STAT-independent, pathway activated in human skin cells that protects against HSV-1 remains to be identified. Non-canonical IFNAR signaling pathways have been described in other cell types(*2, 59*). These include activation of MAPK kinase (MAPK) and phosphoinositide (PI3K)/mammalian target of rapamycin complex (mTOR) pathways, which have transcriptional and non-transcriptional outcomes. However, these non-canonical pathways require JAK/TYK2 activity(*60*), which we found was not required to restrict HSV-1 replication in keratinocytes or organoid fibroblasts. Thus, they are unlikely to contribute to the IFN-κ-mediated HSV-1 restriction detailed herein. Based on our findings, we propose that constitutive IFN-κ production drives the expression of a unique subset of anti-HSV-1 genes through a non-canonical JAK-STAT-independent mechanism. In support of this, we found that some IFN-κ regulated genes (*e.g., C3* and *LCP1*) were not stimulated by recombinant IFN-β treatment of keratinocytes, indicating that IFN-κ may have unique signaling functions compared to the other type I IFNs. Future studies will be necessary to elucidate how these genes are regulated by constitutive IFN-κ production and their contribution to controlling HSV-1 replication in human skin.

Overall, our study highlights the critical antiviral role that constitutive IFN-κ plays in human skin and aligns with recent reports studying the gut and female reproductive tract, where constitutive IFN-λ and IFN-ε expression, respectively, have been implicated in antiviral barrier defense (*61–63*). Whether the IFN-dependent control of virus replication in these barrier tissues occurs through canonical or non-canonical mechanisms have not been thoroughly explored. However, the constitutive production of different IFN family members in distinct barrier tissues may reflect the need to maintain a preexisting antiviral state in tissues that face constant pathogen exposure.

## Acknowledgements

We thank the members of the Orzalli and Rashighi Labs for helpful discussions during the development of this study. Work from this study was supported by the Charles H. Hood Foundation Child Health Research Award (to M.H.O.), the Smith Family Award for Excellence in Biomedical Research from the Richard and Susan Smith Family Foundation (to M.H.O.), NIH grant AI182052 (to M.H.O.) and grants from the DOD (BA230166 and HT9425-24-1-1052) (to K.A.F). M.H.O. holds an Investigators in the Pathogenesis of Infectious Disease Award from the Burroughs Wellcome Fund. J.M.J. was supported by NIH T32 grants AI095213 and GM13575 and F31 AI189083. J.E.G., M.K.S, A.M.C., and J.M.K. are supported by NIH P30 AR075043.

## Materials and Methods

### Primary and immortalized cell culture

Normal human foreskin keratinocytes (NFK) and human foreskin fibroblasts (HFF) were isolated from human foreskin tissue as described previously (*64, 65*) in accordance with University of Massachusetts Chan Medical School IRB protocol #H0021295. Prior to organoid development, NFK were cultured in keratinocyte media (3:1 ratio of Dulbecco’s Modified Eagle Medium (DMEM) low glucose (Gibco, #16000044) and Ham’s F12 (Gibco, #11765054), 1.8 mM adenine, 10 ng/ml cholera toxin, 1x pen/strep, 5% FBS, 80 mM HEPES, 0.25 μg/ml hydrocortisone, 10 ng/ml EGF, and 5 μg/ml insulin) on mitomycin C-treated NIH 3T3 cells (ATCC, #CRL-1658). N/TERT2G keratinocyte cells were cultured in keratinocyte serum-free media (KSFM) supplemented with epidermal growth factor (EGF) and bovine pituitary extract (BPE) from Gibco (#17005042) in accordance with the manufacturer’s instructions. Control and *IFNK^−/−^* N/TERT2G keratinocytes used for organoid development were described previously (*10*). Additional *IFNK*-deficient N/TERT2G keratinocytes used for monoculture experiments were generated by lenti-CRISPR, as described below. Primary and hTERT-HFF(*66*) were cultured in HFF media (DMEM low glucose media (Gibco, #11885076) supplemented with 10% FBS (Gibco, #16000044), 1x Pen/Strep, and 1% 800 mM HEPES. HEK293T, Vero, and U20S cells were cultured in DMEM high glucose media (Gibco, #11965092) supplemented with 5% FBS, 5% BCS (Gibco, #16010159), and 1x Pen/Strep.

### Human skin equivalent and dermal equivalent construction

Human skin equivalents were generated as described previously (*45*) with some modifications. One week before collagen embedding, HFF were thawed and cultured in HFF media. Cells were subcultured four days post-thaw at 1×10^6 cells per 10 cm dish. Two days later, HFF were split 9:1 and replated in 10 cm dishes overnight.

To generate collagen embedded fibroblasts, HFF were lifted by trypsinization and resuspended at 3×10^5^ cells/ml in HFF media. A collagen matrix solution containing 8.5% 10X EMEM (Gibco, #11430030), 0.77% 100X L-glutamine (Gibco, #250300081), 10% FBS, and 7% PureCol type I collagen (advancedbiomatrix, #5005) was pH adjusted using sodium bicarbonate before addition of resuspended HFF. HFF were added to the collagen matrix solution (2.338×10^4^ cells/ml) and mixed with gentle pipetting on ice. The fibroblast containing collagen matrix was dispensed into 6-well inserts (Falcon, #087713) (3 ml/insert) in 6-well deep well plates (Falcon, #08774183) and placed at 37°C for 1 hr to promote collagen polymerization. Following polymerization, 12 ml of HFF media were added to the bottom chamber and 2 ml to the upper chamber of each well. Collagen embedded fibroblasts were maintained at 37°C for 1 week.

NHK or N/TERT2G keratinocytes were thawed the same day as HFF collagen embedding. Keratinocytes were replenished with keratinocyte media (NHK) or KSFM (N/TERT2G cells) 3 days post-thaw. Cells were then split and replated at 2×10^6^ cells per 10cm dish for N/TERT2G cells or at 1×10^6^ cells per 10cm dish with 3×10^6^ mitomycin C-treated 3T3s for NHK. Fresh media (keratinocyte media or KSFM) were added two days after splitting. The following day, keratinocytes were removed from the feeder layer and resuspended at 1×10^7^ cells/ml in epidermalization media I (Epi I; Epi base media [3:1 ratio of DMEM calcium free (Gibco, #11430030) and Ham’s F12, 4 mM L-glutamine, 20 μg/ml adenine (Sigma, #A8626), 0.54 μg/ml hydrocortisone (Sigma, #H-4881), 20 pM triiodothyronine (Sigma, #t5516), 1x insulin transferrin-selenium (ITS-X) (Invitrogen, #51500), 0.1 mM o-phosphorylethanolamine (sigma, #p0503)] supplemented with 628 pg/ml progesterone (Sigma, #p8783), 0.3% chelated bovine calf serum (cBCS) (BCS hyclone, #SH30070), 50 *μ*g/ml sodium ascorbate (NaA) (Sigma, #A4034), 628 pg/ml progesterone (Sigma, p8783), and 10 mg/ml epidermal growth factor (EGF) (peprotech, #AF100-15)). Prior to keratinocyte seeding, media was removed from both the upper and lower compartments of deep well plates containing collagen embedded fibroblasts. Plates were left at room temperature for 20 minutes (with plate lid) to allow the collagen embedded fibroblasts to partially dry. Keratinocytes (50 *μ*l of cell suspension in epi I) were then added dropwise onto the fibroblast containing collagen matrices, ensuring even distribution across the matrix surface. Collagen matrices were left undisrupted at room temperature for 15 minutes to allow keratinocyte attachment and were subsequently incubated at 37°C for 1 hr. After incubation, 12 ml of Epi I media were added to the bottom and 2 ml in the upper chamber of the 6-well plates.

Three days after keratinocyte seeding, media was removed from both chambers of 6-well deep well plates. Epidermalization media II (Epi II) (Epi base media supplemented with 0.3% BCS, 50 *μ*g/ml NaA, 628 pg/ml progesterone, and 132.5 mg/ml calcium chloride [CaCl_2_] [Sigma, C7902]) was then added to each well (12 ml to the lower chamber and 2 ml to the upper chamber). Two days after addition of Epi II, media was removed from both chambers and 10 ml of cornification media (Cornification base media [1:1 ratio of calcium free DMEM and Ham’s F12, 4 mM L-glutamine, 24 μg/ml adenine, 0.5 μg/ml hydrocortisone, 20 pM triiodothyronine, 1x ITS-X, 0.1 mM O-phos], supplemented with 2% BCS, 50 μg/ml NaA, 265 μg/ml CaCl2) was added to the bottom chamber establishing an air liquid interface. Organoids received fresh cornification media on the second day post-media change and were ready for experimentation on day 5.

Dermal equivalent organoids were generated using the same procedure described above but without keratinocyte seeding.

### Viruses and viral infections

The HSV-1 WT strain KOS (*67*) and HSV-1 K26-GFP (*47*) were grown and viral titers were determind by plaque assay on Vero cells following standard procedures (*36*). HSV-1 ΔICP0 (7134) and HSV-1 ΔICP0 (7134R) (*68*) were grown and titers defined on U2OS cells. VSV (*69*) was propagated on BHK-21 cells and titers defined on Vero cells. Sindbis virus p389-GFP (SINV-GFP) and Venezuelan equine encephalitis virus (VEEV-TC-83-GFP) were generated as previously described (*70, 71*). Infectious viral stocks were generated by electroporation of *in vitro* transcribed viral RNA into BHK-21 cells. At 36–48 h post-electroporation, supernatants were collected and stored at −80°C as passage 0 (P0) stocks. High-titer P1 stocks were subsequently generated in BHK-21 or Vero cells, and viral titers were determined by plaque assay.

For WT HSV-1 KOS, HSV-1 ΔICP0, HSV-1 ΔICP0R, and VSV infections, keratinocytes were seeded in 24 well plates (3×10^5^ cells per well) 24 hrs prior to infection. Viruses were diluted to the indicated MOI in KSFM, added to cells, and incubated at 37°C for 1 hr with gentle shaking. After 1 hr virus absorption, virus containing media was aspirated from cells and replaced with KSFM. For SINV 389-GFP and VEEV-TC-83-GFP infections, keratinocytes were seeded in 96-well plates (1×10⁴ cells/well) and infected at the indicated MOI. At 48 hpi, cells were fixed with 2% paraformaldehyde (PFA) and stained with Hoechst for 10 minutes. GFP fluorescence was measured using a Cytation 5 imaging reader.

DE and HSE organoids were infected with 1×10^4^ PFU of WT HSV-1 KOS, HSV-1 K26-GFP, or VSV. Organoids were first removed from 6-well inserts and transferred to a 10 cm dish. The air exposed surface of the organoids was treated with 100 *μ*l of dispase II solution (2.5 mg/ml) (Sigma, 4942978001) for 2 minutes. Following dispase II treatment, the epiderms from HSE organoids was mechanically removed with tweezers. The fibroblast embedded collagen matrix was then transferred to 6-well plates with the non-dispase II-exposed surface facing up. Virus was diluted in 100 *μ*l of PBS (Gibco, 14190250) and applied to the surface of each organoid. Following a 1 hr incubation at 37°C, 2 ml of cornification media was added to each well. Organoid samples were collected at the indicated time points for downstream analysis.

### Chemical inhibitors

Abrocitinib (#HY-107429, MCE) and Deucravacitinib (#HY-117287, MCE) were resuspended in DMSO and added to N/TERT2G keratinocytes and organoid cultures 24 hr prior to infection at a final concentration of 10 *μ*M. After 1 hr of infection, KSFM (for keratinocytes) or cornification media (for organoid cultures) containing 10 *μ*M of Abrocitinib and Deucravacitinib was used to replace virus containing media.

### Immunoblots

Cells were lysed in 1x Laemmli buffer containing β-mercaptoethanol and incubated on a 98°C heat block for 5 minutes. Samples were run on 10% tris-glycine gels and transferred to immobilon-P PVDF membranes overnight at 4°C. Membranes were blocked for 30 min in tris-buffered saline with 0.05% tween (TBST) containing 5% milk followed with overnight incubation in primary antibodies diluted in TBST containing 2.5% bovine serum albumin (TBST-BSA). Blots were washed 3x in TBST for 10 min and incubated with goat anti-rabbit or goat anti-mouse secondary antibodies diluted in TBST with 5% milk at room temp for 1 hr. Membranes were then washed 2x in TBST followed by 1x wash in TBS for 10 minutes each. Blots were developed using Pico or Femto supersignal western blot substrate (Thermo fisher scientific) and imaged on a Chemidoc MP imaging system (Biorad). Primary antibodies used for immunoblots were IFN-κ (Abnova, #H00056832-MO1), STAT1 (cell signaling, #9172S), P-STAT1(BD Biosciences, #612233), beta-Tubulin (cell signaling, #86298S), IRF9 (cell signaling, #76684T), STAT2 (cell signaling, #72604), IFNAR2 (cell signaling, #53883T), and IFNAR1 (proteintech, #83002-4-RR).

### Plaque Assays

For HSV-1 plaque assays, lysates from infected keratinocytes were collected at the indicated timepoints by adding an equal volume of autoclaved 10% milk (Nestle Carnation instant dry milk) diluted in water. Sample lysates were frozen at −80°C and freeze thawed twice on ice. Thawed samples were serially diluted (10-fold dilutions) in DMEV (DMEM supplemented with 1% FBS and Pen/strep) and added to a monolayer of Vero (HSV-1 KOS) or U2OS cells (HSV-1 ΔICP0 and HSV-1 ΔICP0R) in a 6-well dish, then incubated at 37°C for 1 hr with shaking. The virus inoculum was then aspirated, and cells were overlayed with DMEV supplemented with human immunoglobulin (IgG) at a concentration of 10 μg/ml (Sigma-Aldrich, #I44506). 72 hours post-infection, DMEV containing IgG was removed and cells were fixed with ice-cold 100% methanol and then stained with crystal violet resuspended in 10% methanol. Plates were air dried overnight and viral titer was determined by counting plaques.

For VSV plaque assays, supernatants from VSV infected cells or organoids were collected at the indicated time points and frozen at 80°C. Supernatant samples were thawed on ice and serially diluted (10-fold dilutions) in DMEV and added to a monolayer of Vero cells in 6-well dishes, then incubated at 37°C for 1 hr with shaking. Virus inoculum was aspirated and cells were overlayed with 1x MEM (Fisher, #61100087) containing 0.25% sterile agarose. 18 hours post-infection cells were fixed with 10% formalin for 1 hr at RT. Agarose overlays were then removed and cells were stained with crystal violet resuspended in 10% methanol. Plates were air dried overnight and viral titer was determined by counting plaques.

### Flow cytometry

Collagen embedded fibroblasts were incubated in a collagenase A (Roche, #10103586001) containing buffer at 37°C with periodic shaking until collagen dissolved (∼1 hr). Remaining fibroblasts were fixed in 4% paraformaldehyde on ice for 10 minutes. After fixation, fibroblasts were resuspended in PBS. Cells were analyzed using the Biorad ZE5 Fluorochrome. Acquired flow cytometry data was analyzed with FlowJo software.

### RNA isolation and RT-qPCR

RNA was isolated from collagen embedded HSE and DE fibroblasts using the NEB Total RNA miniprep kit and eluted in 50 μl of nuclease free water. For detection steps, a total of 120 ng of RNA was used. Reverse transcription and qPCR reactions were performed using the Taqman RNA-to-CT 1-Step kit following the manufacturer’s instructions. Taqman primer probes used for gene detection *MX2* (Hs01550814)*, BST2 (*Hs00171632), *IFITM1*(Hs00705137), and *TBP* (Hs00427620) were acquired from Thermofisher.

### Bulk RNA-seq analysis

Total RNA was isolated from Cas9-control and *IFNK*-deficient N/TERT2G cells from three independent biological replicates per condition. Samples with RNA integrity numbers (RIN)> 8 were used for library preparation. RNA-seq libraries were prepared and sequenced by Azenta Life sciences to generate paired end reads with ∼ 20M reads per sample.

For IFN-β treated keratinocytes, N/TERT2G keratinocytes were seeded at 3×10^5^ cells per well in 24-well plates. The next day cells were treated with recombinant IFN-β (peproTech, 300-02BC-5UG) at 1 ng/ml or vehicle control (DMSO) for 6 hr, with three biological replicates per condition. RNA-seq libraries were generated and sequenced by Plasmidsaurus using a 3’ end counting approach on an Illumina platform yielding ∼11M reads per sample.

For experiments with DE and HSE organoids, total RNA was isolated as described above for qRT-PCR analysis and subjected to poly-A selection. RNA-Seq libraries were prepared and sequenced by Azenta Life Sciences to generate paired-end reads (2×150bp) with a depth of ∼20M reads per sample.

The ViaFoundry RNA-seq pipeline (v2.7.4) was used for quality control, genome alignment and estimating gene expression counts (*72*). Raw sequencing reads were quality-checked using FastQC and aligned to the human reference genome (hg38, v44) using STAR. RSEM was used to estimate gene expression levels. Differential gene expression analysis was performed using DEBrowser (v1.0.0) in R. Genes with at least 5 counts per million in at least half the samples were retained for analysis. Differentially expressed genes were defined using a threshold of −1.0 < log_2_FC >1.0 and padj < 0.05. Gene set enrichment analysis (GSEA) was performed using fgsea R package (v1.36.2). Heatmaps were generated using clusterprofiler (v4.18.4), and volcano plots were generated using ggplot2 (v4.0.2).

### CRISPR gene targeting

Guide RNAs targeting *IFNK (*sg*IFNK* #3: GCAGCAATGAATATACCCATA, sg*IFNK* #4: GTTTGTGGCTTGAGATCCTTA)*, IFNB1 (*sg*IFNB*1 #1: GTCCAAGCAAGTTGTAGCTCA, sg*IFNB1* #3: GCTCATGGAAAGAGCTGTAG), *IFNAR1 (*sg*IFNAR1* #3: GTAGATGACAACTTTATCCTG, sg*IFNAR1* #5: GTTCCATCAGATGCTTGTACG)*, IFNAR2 (*sg*IFNAR2* #1: GAGGTAACATGGTTGAGTGA)*, STAT1 (*sg*STAT1* #1: GCATGCCAACGCACCCTCAG)*, STAT2 (*sg*STAT2* #3: GCAAGAAGTGGAAGAATAGCA), and *IRF9 (*sg*IRF9* #1: GCTTTCCCTTAAATATTGCCC), were designed using synthego.com. Double stranded DNA oligos containing guide RNA sequences of interest were cloned into AfeI/SbfI digested PRRL-Cas9 PURO lentivirus construct (*73*). Lentivirus constructs (6 μg) were transfected into HEK293T cells along with the plasmids pVSV-G (1.5 μg) and psPAX2 (3 μg) using polyethylenimine (40 μl, stock 1 μg/ml) (#408727; Sigma-Aldrich). Twenty-four hours after transfection, transfection complexes were replaced with HFF media or DMEM supplemented with 10% FBS and 1x pen/strep for samples to be used for fibroblast or keratinocyte transduction, respectively. For fibroblasts, supernatants from the transfected HEK293T cells were collected at 24 hours post-transfection, filtered through 0.45 μm filters and added to 6-well dishes containing primary human fibroblasts plated at 1×10^5 cells per well. Fibroblasts were transduced by spinfection at 2,000 RPM for 30 minutes. Forty-eight hours post transduction, lentivirus containing media on fibroblasts was replaced with HFF media containing puromycin (1 μg/ml) for selection until all control cells were dead.

For N/TERT2G keratinocytes, HEK293T cell supernatants were collected at 48 hours post-transfection and lentivirus was concentrated using the Lenti-X concentrator (#631231; Takara Bio). Lentivirus pellets were resuspended in KSFM, added to N/TERT2G keratinocytes, and transduced by spinfection at 2,000 RPM for 30 minutes. Twenty-four hours after transduction, N/TERT2G keratinocytes were cultured in KSFM with puromycin (20 μg/ml) until all control cells were dead. Single cell clones for N/TERT2G cell lines were generated by plating limiting dilutions of cells in 10 cm dishes (100-200 cells per plate). Single cell colonies were isolated using 8 mm cloning cylinders (Corning, 3166-8) and autoclaved dow corning high vacuum grease (Dupont, #3173528).

### scRNA-seq analysis

Data from a previous publication (GSE179633) was analyzed as described in (*74*). Samples from healthy skin were visualized and exported from https://CD14photosensitivity.umassmed.edu

### Statistical Analysis

GraphPad Prism v11 was used for all statistical analysis noted in the figure legends.

### Data availability

Raw and processed RNA-seq data is deposited at NCBI Gene Expression Omnibus (GSE343241, GSE343242, GSE343459, GSE343460).

---

**Figure S1.**
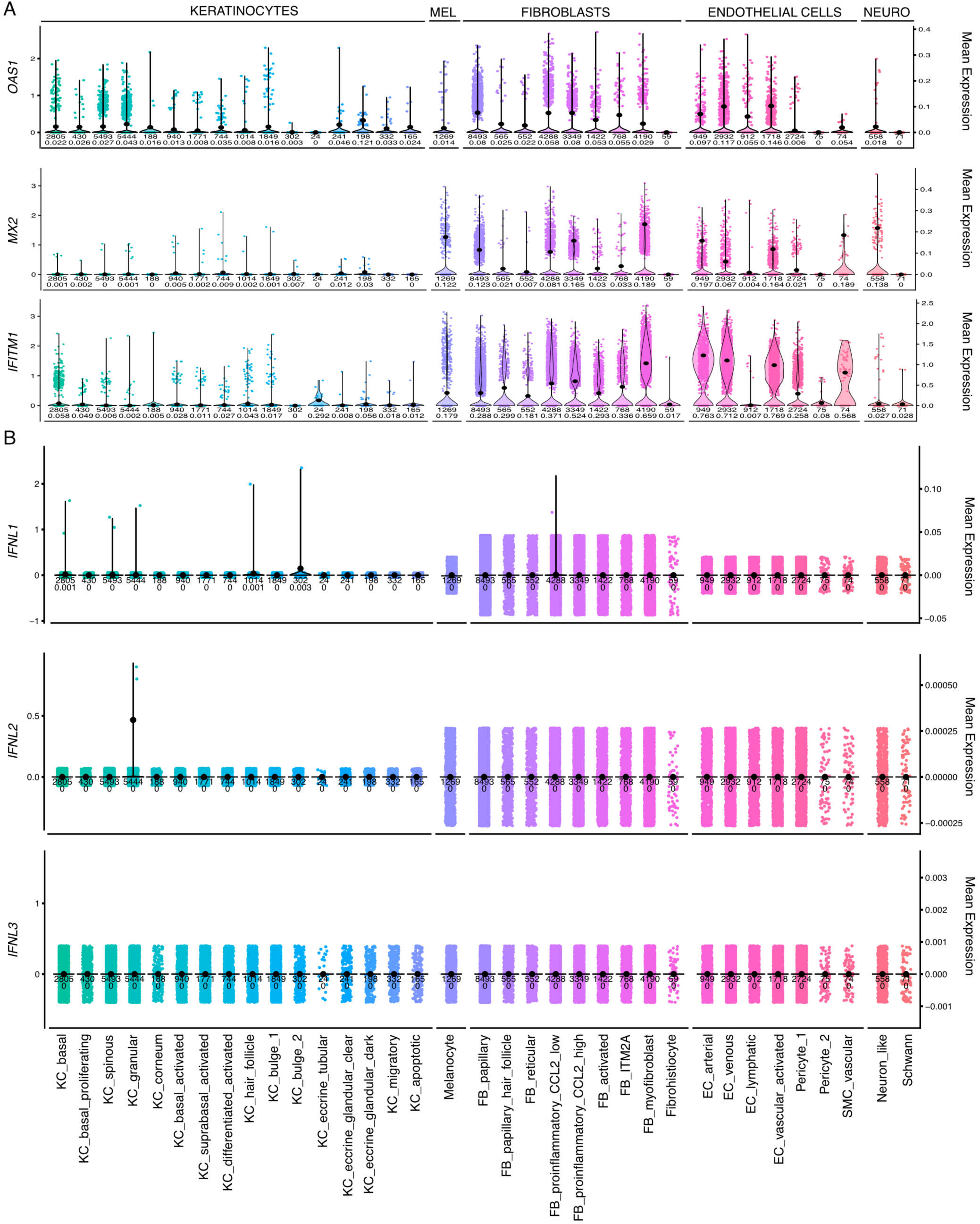
Antiviral transcripts are detected in healthy human skin cells. A) Violin plots showing expression of the ISGs *OAS1*, *MX2*, and *IFITM1* in keratinocyte, melanocyte, fibroblast, endothelial cell, and neuronal cell subtypes from healthy skin punch biopsies (GSE179633). Each dot represents a single cell. B) Violin plots showing expression of IFNλ subtypes in keratinocyte, melanocyte, fibroblast, endothelial cell, and neuronal cell subtypes from healthy skin punch biopsies (GSE179633). Each dot represents a single cell.

**Figure S2.**
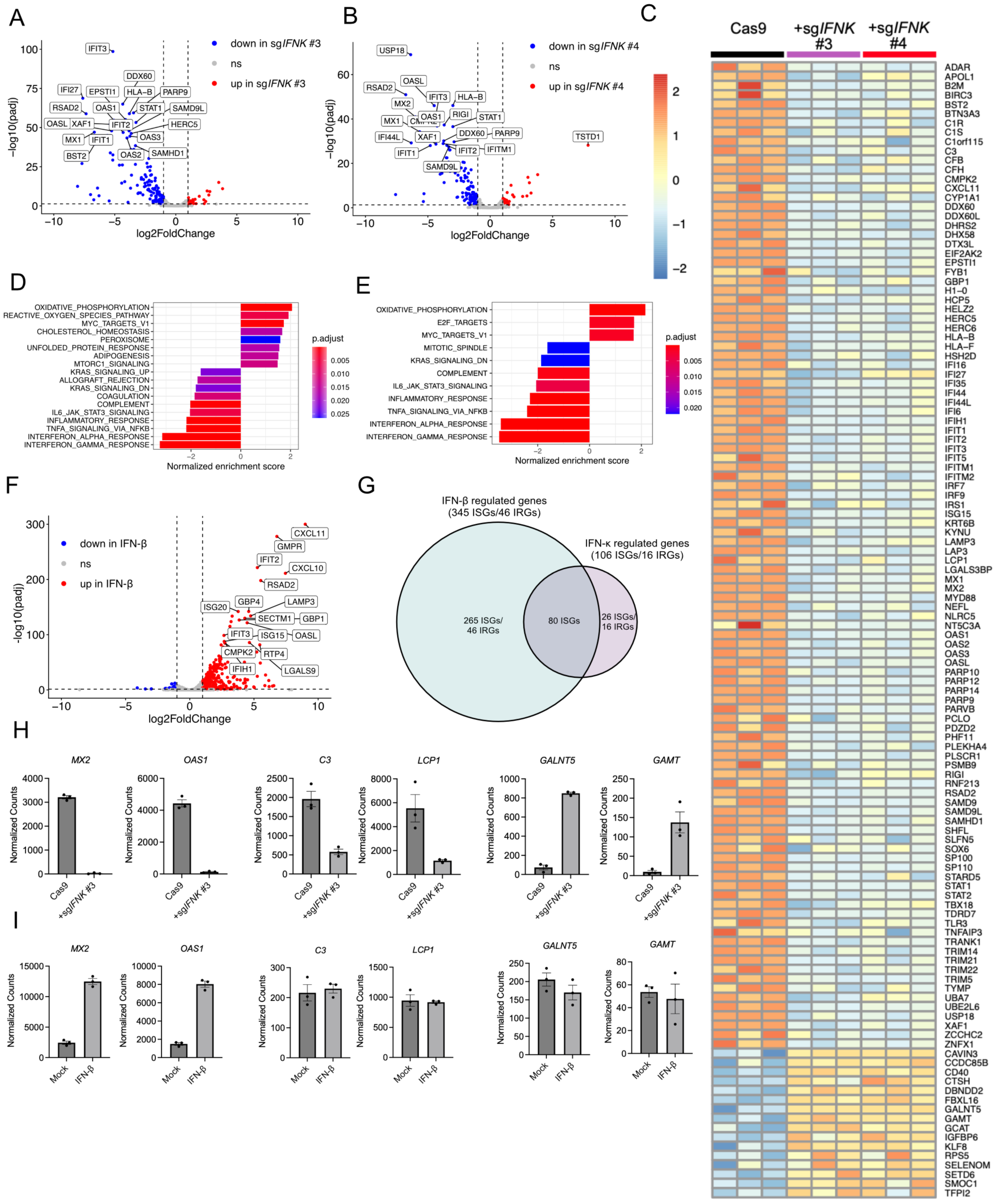
*IFNK*-regulates expression of canonical and non-canonical ISGs in keratinocytes. A and B) Volcano plots showing differentially expressed genes in A) sg*IFNK* #3 and B) sg*IFNK* #4 N/TERT2G keratinocytes compared to control (Cas9) cells. Dashed lines indicate differentially regulated genes with a cut off of −1.0 < log2FC >1.0 and padj < 0.05. C) Heat map of RNA-seq data showing all differentially expressed genes that are shared between sg*IFNK* #3 and sg*IFNK* #4 N/TERT2G keratinocytes compared to control (Cas9) cells. D and E) GSEA analysis of D) sg*IFNK*#3 and E) sg*IFNK* #4 N/TERT2G keratinocytes compared to control cells. F) Volcano plot showing differentially expressed genes from N/TERT2G cells treated with recombinant IFN-β (1 ng/ml) for 6 hrs. Dashed lines indicate differentially regulated genes with a cut off of −1.0 < log2FC >1.0 and padj < 0.05. G) Venn diagram showing overlap between recombinant IFN-β and IFN-*κ* regulated genes in N/TERT2G keratinocytes. H) Normalized counts for *MX2*, *OAS1*, *C3*, *LCP1*, *GALNT5*, and *GAMT* transcripts from RNA-seq data from control (Cas9) and sg*IFNK* #3 N/TERT2G cells. I) Normalized transcript counts for *MX2*, *OAS1*, *C3*, *LCP1*, *GALNT5*, and *GAMT* transcripts from RNA-seq data from N/TERT2G cells treated with recombinant IFN-*β* (1 ng/ml) for 6 hrs.

**Figure S3.**
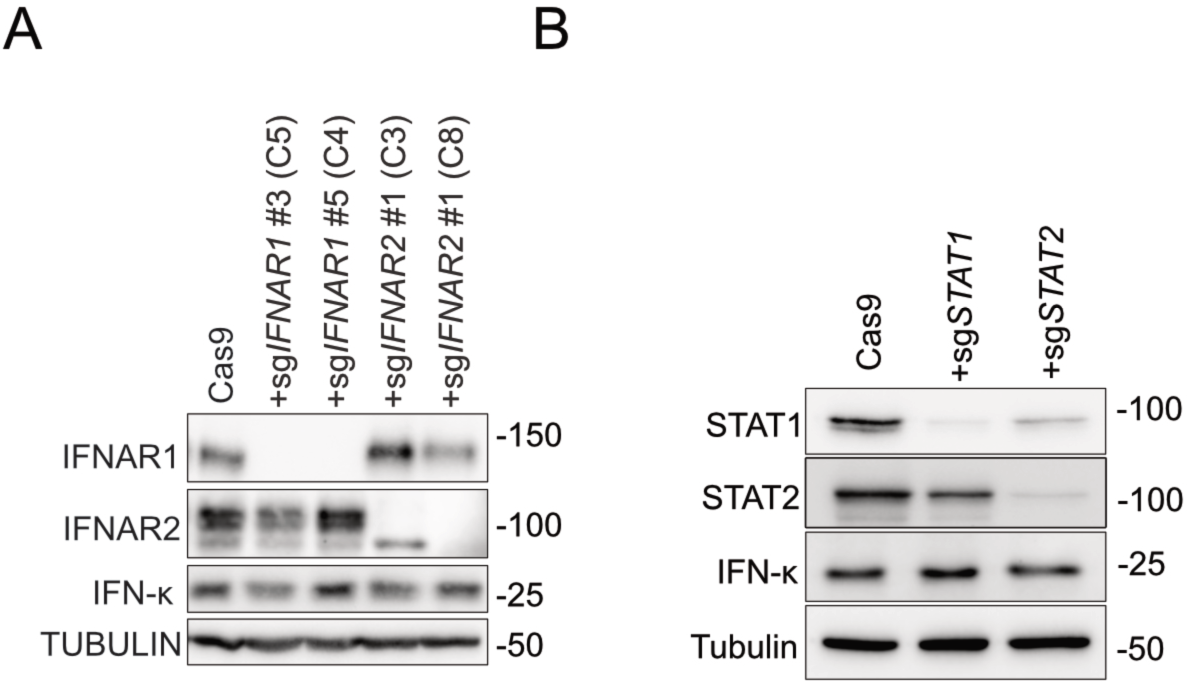
Validation of IFNAR1/2 and STAT1/2-deficient keratinocytes. A) Immunoblots of lysates from single cell clones (C#) from control (Cas9), *IFNAR1*-deficient, and *IFNAR2*-deficient N/TERT2G keratinocytes showing IFNAR1, IFNAR2, and IFN-*κ* protein abundance. Images are representative of n=3 biological replicates. B) Immunoblots of Cas9, *STAT1-*, and *STAT2-*deficient N/TERT2G keratinocytes showing STAT1, STAT2, and IFN-*κ* protein abundance. Images are representative of n=3 biological replicates.

**Figure S4.**
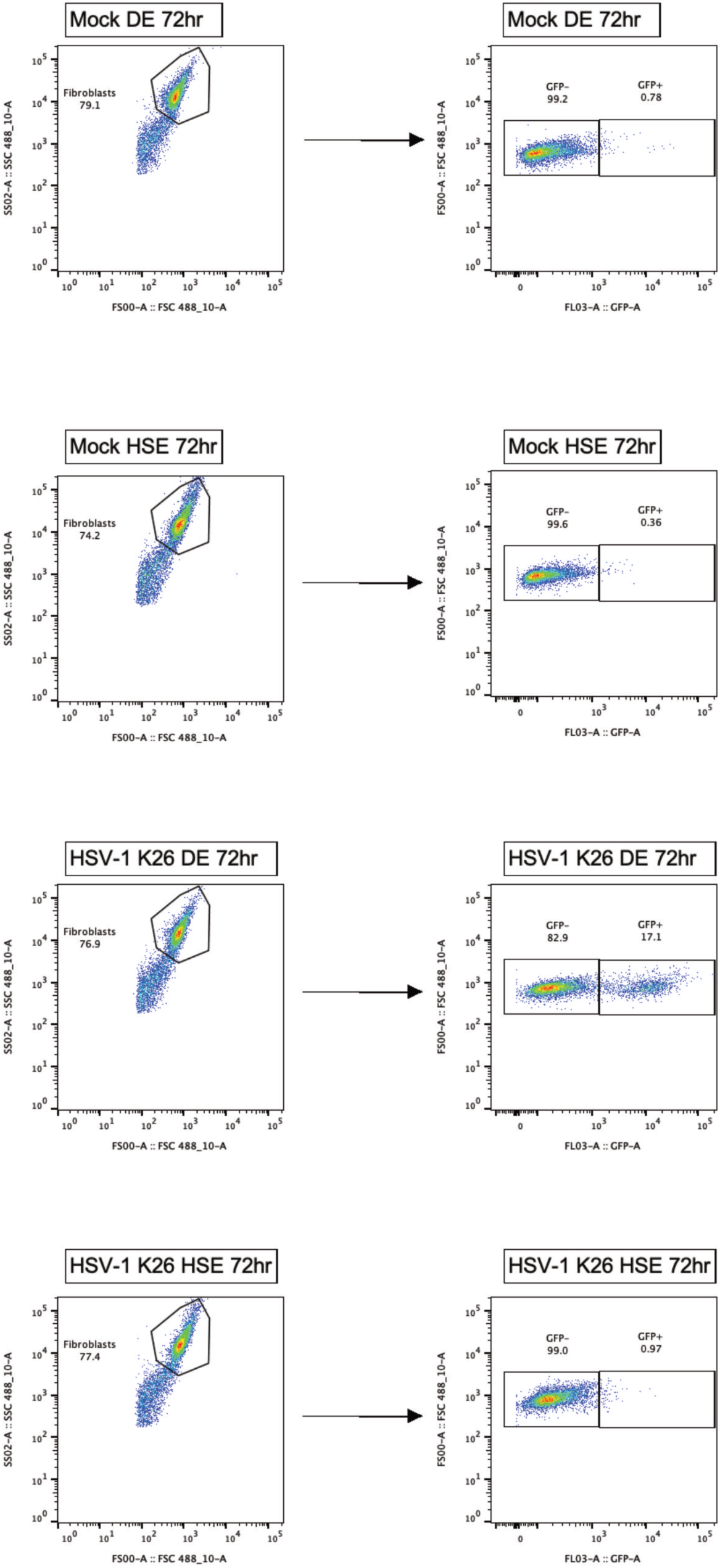
Representative flow analysis of HSV-1 K26-GFP infected organoid fibroblasts Gating strategy and representative flow plots used to identify GFP+ fibroblasts from HSV-1 K26-GFP-infected organoid cultures.

**Figure S5.**
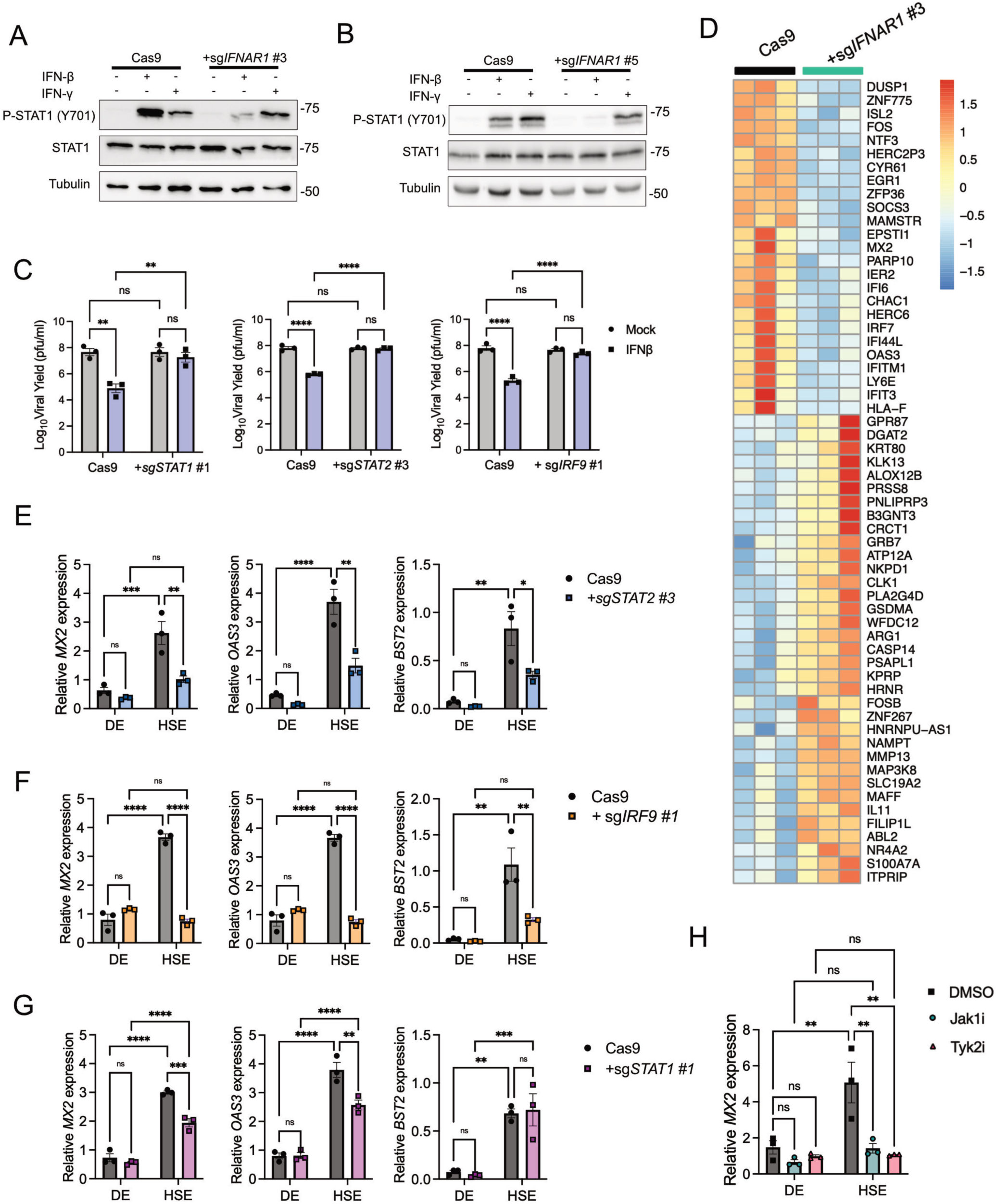
Extended data to support Figure 6. A and B) Immunoblots of lysates from Cas9 and *IFNAR1*-deficient fibroblasts treated with recombinant IFN-β or IFN-ɣ (100 pg/ml) for 15 minutes showing p-STAT1 (Y701) and total STAT1 protein abundance. C) Comparison of viral yields in Cas9, *STAT1*-, *STAT2*-, and *IRF9*-deficient fibroblasts infected with VSV (MOI 1.0) for 24 hrs and treated with IFN-β (100 pg/ml) at 1 hpi. Data are presented as the mean ± SEM of n=3 independent experiments. D) Heat map representation of RNA-seq data showing differentially expressed genes in Cas9 and *IFNAR1*-deficient HSE fibroblasts. DEseq2 was used to identify differentially expressed genes with a −1.0 < log2FC >1.0 and padj < 0.05 cut off. E-G) *MX2* transcript abundance relative to *TBP* in DE and HSE organoid fibroblasts generated with G)*STAT1*-, F) *STAT2*-, and E) *IRF9*-deficient fibroblasts compared to Cas9 control cells. Data are presented as the mean ± SEM of n=3 independent experiments. H) qRT-PCR analysis of *MX2* transcript abundance relative to *TBP* in DE and HSE fibroblasts treated with 50 μM of Abrocitinib (JAK1i) or Deucravacitinib (TYK2i) for 24 hrs. Data are presented as the mean ± SEM of n=3 independent experiments.

**Figure S6:**
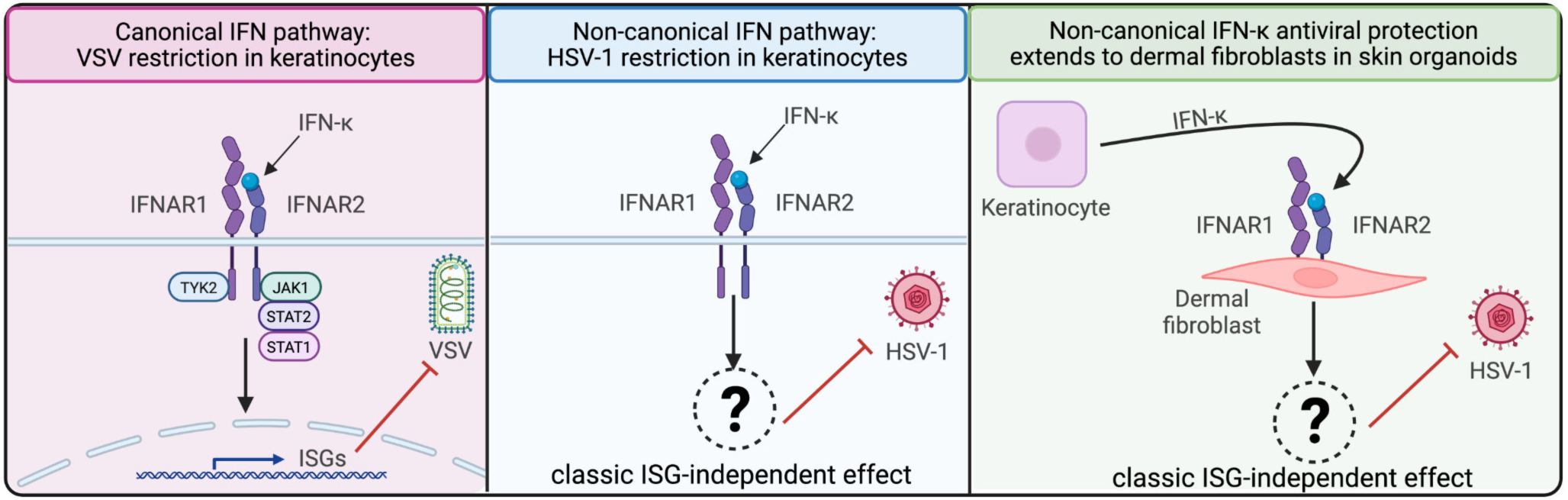
Model figure for IFN-κ-mediated establishment of a pre-existing antiviral state in human skin.

